# CPAP/CENPJ is essential for the stability and function of ESCRT-pathway associated AAA+ ATPase VPS4B

**DOI:** 10.64898/2026.05.29.728888

**Authors:** Radhika Gudi, Mousumi Bhattacharjee, Chenthamarakshan Vasu

## Abstract

Function of CENPJ/CPAP is essential for centriole duplication and cilia biogenesis, disruption of which can lead to microcephaly and Seckel syndrome. Recently, we showed that CPAP is an integral Endosomal Sorting Complexes Required for Transport (ESCRT)-0-like protein that recruits ESCRT-I protein TSG101 to early endosome (EE) and it positively regulates multi-vesicular body (MVB) formation. Sequential recruitment of the ESCRT protein complexes and AAA+ ATPase VPS4B to EE facilitates MVB biogenesis. VPS4B is critical for ESCRT-III disassembly/recycling and contributes to membrane fission in several cellular processes. Here, we report that CPAP is critical for the protein stability and EE localization of VPS4B, and this function is independent from its role as an ESCRT-0 protein. Other well-recognized VPS4B-dependent cellular processes such as cytokinesis, extracellular vesicle production, and retroviral budding are also compromised under CPAP deficiency. We found that CPAP overexpression leads to cellular accumulation and CPAP deficiency causes proteasomal degradation of VPS4B. We also found that VPS4B interacts with the C-terminal CC4 domain of CPAP, which is considered as its centriole localization domain, and this interaction prevents the proteasome degradation of VPS4B. Notably, the EE localization of VPS4B can be attributed to microcephaly-associated C-terminal TCP domain of CPAP. Overall, these observations provide evidence that CPAP is critical for VPS4B and its ESCRT-associated function and suggest that distinct pools of CPAP may be involved in its ESCRT-0 and VPS4B stabilization roles. This study also provides clues to potential mechanisms underlying the clinical phenotypes of CPAP deficiency.

## Introduction

Endocytosis is a fundamental biological process that maintains the cellular, tissue and organismal homeostasis. Post endocytosis, cell receptor cargo traffics through, and is sorted by, a series of tubulovesicular compartments referred to as endocytic vesicles (1). Receptors such as epidermal growth factor receptor (EGFR) are routed through different functional stages of endosomes and ultimately reach the lysosome for degradation. Early endosomes (EE) serve as a sorting nexus that decides the fate of endocytosed cargo (1–3). However, formation of the multi-vesicular body (MVB), which is composed of a limiting membrane that encloses several smaller vesicles carrying the cargo (referred to as intraluminal vesicles [ILVs]), is an essential step during cargo sorting (4,5). Inward budding and scission of cargo-associated membrane to generate ILVs that are enclosed by the limiting membrane of MVBs requires the sequential recruitment of ESCRT protein complexes (0, I, II, III and the accessory counterparts including AAA-type ATPases; paralogs VPS4A and VPS4B) to the EE membrane (6–9). The early acting ESCRT proteins recruit subunits of ESCRT-III complex, resulting in the formation of filaments around the necks of membrane tubules. These ESCRT-III assemblies, particularly late players CHMP2A and CHMP3, in turn, recruit VPS4 ATPases. ATP hydrolysis by VPS4 causes the scission of ILVs by orchestrating disassembly/reassembly cycles of the ESCRT III complexes (10–13). Mature MVB or late endosomes are fused with lysosomes to complete the endocytic vesicular transport (EVT) of cargo proteins targeted for degradation. The ATPase function of VPS4B is also critical for other cellular processes such as exosome biogenesis, cytokinesis, and viral budding (14–16).

Recently, we reported a novel ESCRT-associated positive regulatory function of CPAP (Centrosomal P4.1-associated protein; expressed by CENPJ gene)/SAS-4, a microtubule/tubulin-binding essential centriole biogenesis protein (17–22) on EVT of internalized EGFR cargo and endosome maturation (23,24). Employing gain-and loss-of-function approaches as well as the ligand-bound EGFR intracellular-trafficking model, we demonstrated that CPAP is required for the efficient transport of internalized EGFR to MVB and lysosome for degradation (23). Importantly, higher EGFR function due to defective EVT function under CPAP-deficiency elevated the epithelial-mesenchymal transition (EMT) and increased the tumorigenicity of oral cancer cells (25). We also showed that CPAP functions in parallel to HRS and as an ESCRT-0-like protein and recruitment of a key ESCRT-I protein, TSG101 to EE is one of the molecular mechanisms by which CPAP promotes MVB formation and endosome maturation (24).

Here, we report a surprising, CPAP-associated molecular event, far downstream of ESCRT-I recruitment by which MVB formation and endosomal trafficking of receptor cargo are facilitated. We show that the cellular levels of VPS4 proteins and endosome localization of VPS4B paralog are profoundly diminished in CPAP-depleted cells and enhanced in CPAP-overexpressing cells. In addition to the defective MVB formation and lysosomal targeting of EGFR (23,24), we found that CPAP deficiency negatively impacted other major VPS4B-dependent cellular events such as cytokinesis, exosome production, and viral egress. We observed that CPAP interacts with, and stabilizes, VPS4B through a well-defined centriole localization domain in its C-terminus (CC4 domain) (26). However, the endosome localization of VPS4B also requires another C-terminal domain, STIL-binding structural-motif called TCP (26,27), in CPAP which is known to be critical for proper centriole formation and avoiding microcephaly (26,28). Overall, these studies suggest that complex formation with CPAP is required for VPS4B protein stability and thereby its ATPase function, which is essential for efficient ILV budding and MVB formation, as well as other obligate VPS4B-dependent cellular events including exosome biogenesis, cytokinesis, and viral budding. Our current observations re-emphasize that CPAP is an essential component of ESCRT pathway and, perhaps distinct pools of, this protein could function in parallel to facilitate multiple cellular events.

## Results

### Cellular levels of VPS4 proteins are severely diminished upon CPAP depletion

Our recent report (24) showed that CPAP functions as an ESCRT-0 protein and recruits ESCRT-I protein TSG101 on to endosome, without affecting the cellular levels of HRS, TSG101 or ALIX. Our reports (23,24) have shown that endosome maturation and MVB formation are disrupted in CPAP-depleted cells. Hence, by employing an IB assay, we examined whether CPAP depletion impacts the cellular levels of critical ESCRT pathway-associated AAA^+^ ATPases, VPS4A and VPS4B, which are known to be essential for ILV scission and MVB formation. Of note, human VPS4A and VPS4B paralogs are often listed as VPS4 in the literature, hence the antibodies used in our study were validated using VPS4A-and VPS4B-depleted cells **(Fig. S1A)**. We found that cellular levels of VPS4 proteins are significantly diminished in HEK293T and HeLa cells when CPAP is depleted using a custom pool of four CPAP-specific siRNAs **(Fig. 1A)** suggesting that CPAP is required for the stability of VPS4 proteins. To rule out the non-specific effects on VPS4, if any upon using multiple pooled siRNAs, individual CPAP-specific siRNAs were also tested for depleting CPAP and the impact on VPS4 levels. As observed in **Fig. S1B**, CPAP depletion using all four siRNAs individually resulted in diminished cellular levels of VPS4 proteins. Of note, similar reduction in VPS4 protein levels was observed in spleen and primary fibroblasts from CPAP hypomorphic mice **(Fig. S2)** suggesting that genetic depletion of CENPJ in mice also causes reduced VPS4B protein levels and CPAP protein is required for VPS4 stability in mice and humans. Further, as shown in **Fig. S3A**, depletion of VPS4A or VPS4B using specific siRNAs had no impact on cellular levels of CPAP protein suggesting that CPAP is required for VPS4 stability, but not vice versa. Possible off target effect of the pool of CPAP specific siRNAs on mRNA of ESCRT-associated proteins was also assessed by quantitative RT-PCR (qRT-PCR). As observed in **Fig. 1B**, cells treated with CPAP-specific siRNA showed diminished mRNA levels of CPAP, but not that of ESCRT-associated proteins VPS4B, VPS4A, HRS, TSG101 and ALIX. In fact, an increase in the mRNA levels of these ESCRT associated proteins was observed in CPAP-siRNA treated cells as compared to control siRNA-treated cells possibly due to CPAP deficiency-associated general cellular stress. This, in the context of our previous report showing CPAP levels do not influence HRS, VPS4B and ALIX protein levels (24), suggest that CPAP deficiency selectively destabilizes cellular levels of VPS4 proteins.

**Fig. 1.**
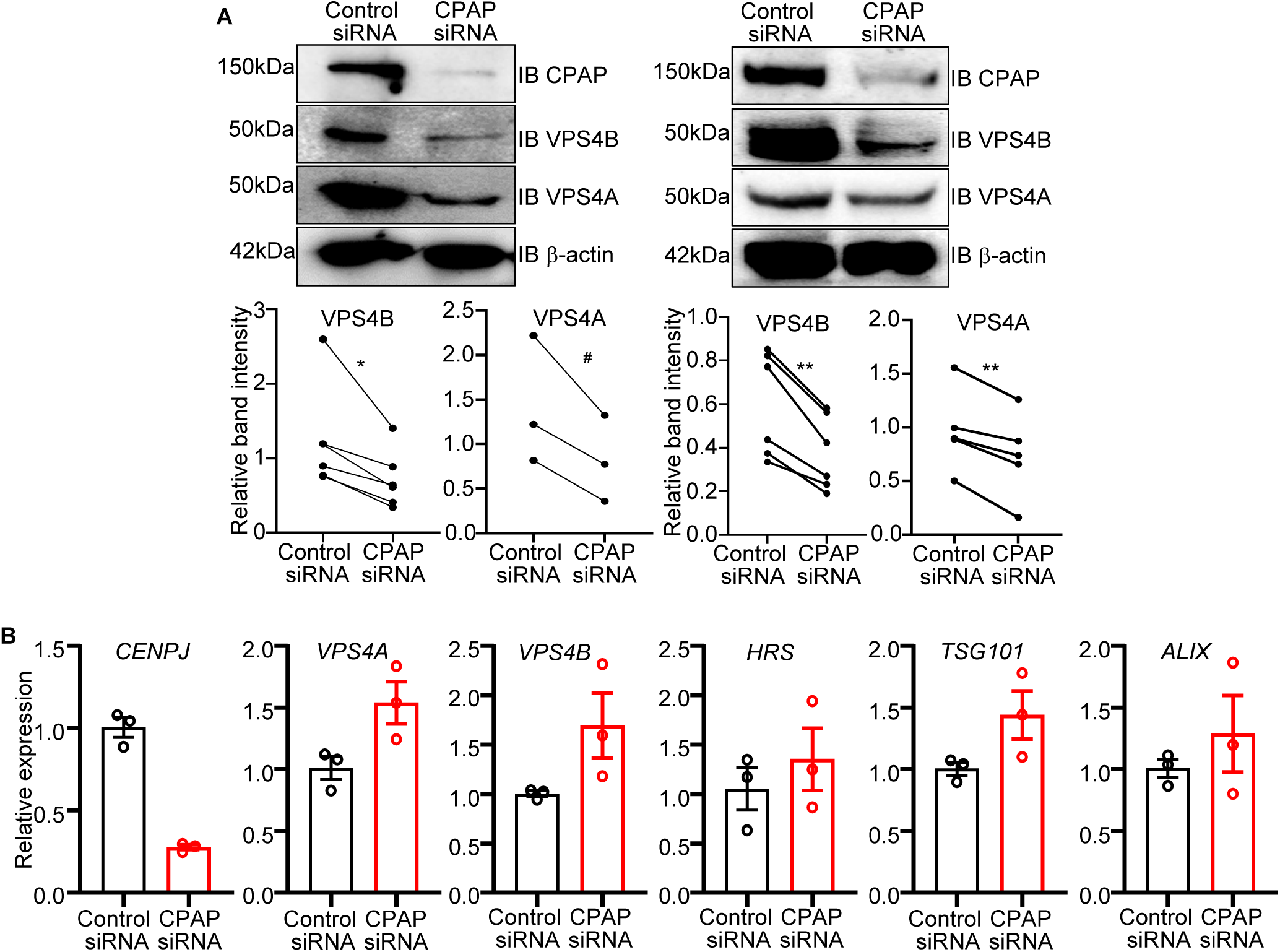
Cellular levels of VPS4 proteins are severely diminished upon CPAP depletion. **(A)** HEK293T (left panel) or HeLa (right panel) cells were treated with scrambled-control or a pool of four CPAP-specific siRNA for 48h and subjected to IB to detect the cellular levels of VPS4B and VPS4A proteins. Representative blots (upper panels) and densitometric values of VPS4 protein bands (relative to β-actin) from at least three independent experiments (lower panels), are shown. **(B)** cDNA prepared from HEK293T cells that were treated with scrambled-control or a pool of four CPAP-specific siRNA for 24h were subjected to qPCR to detect the mRNA levels of CPAP and ESCRT-associated proteins. Specific factor cDNA levels relative to β-actin levels were calculated first and the relative expression in CPAP-depleted cells was calculated against control siRNA-treated cells. Mean ± SD values of cells from three parallel cultures are shown. *p*-values by paired *t*-test for panel A and Mann-Whitney test for panel B. ^#^=0.054, *<0.05, **<0.01.

It has been reported that endocytosis of lysosome-targeted cell surface receptors leads to the sequential recruitment of ESCRT protein complexes and accessory proteins, including VPS4A and VPS4B, to the EE. ESCRT-III filaments constrict and cut the membrane neck to form ILVs (13) and the VPS4 proteins promote the dissociation/reassociation of ESCRT-III complex until membrane scission is achieved (29). Since our recent report found that CPAP exerts ESCRT-0 function (24), we examined if depletion of early acting ESCRT proteins HRS (ESCRT-0) or TSG101 (ESCRT-I) produces similar effects to that of CPAP-depletion on VPS4 protein levels. **Fig. S3B** shows that depletion of HRS or TSG101 proteins does not affect the cellular levels of VPS4B and VPS4A. These results also suggest that the effect of CPAP deficiency on cellular stability of VPS4 proteins, the downstream ESCRT-associated ATPases that are generally considered to be recruited by ESCRT-III, is possibly independent of CPAP’s ESCRT-0 function that we reported recently (24).

### Endosome localization of VPS4B is disrupted in CPAP depleted cells

We then examined if the EE recruitments of VPS4A and VPS4B proteins are impacted in CPAP-depleted cells undergoing EVT. EGF-treated control and CPAP-depleted HeLa cells were stained for VPS4A and VPS4B, using antibodies that are validated by IF staining approach **(Fig. S4)**, together with the anti-EEA1 antibody (to mark EE) and subjected to Airyscan super-resolution microscopy. We found that the recruitment of VPS4B to the EEA1-positive vesicular structures, as assessed by quantifying the percentage of EEA1 puncta positive for VPS4B puncta in single Z scan images, was significantly lower in CPAP-depleted cells compared to control cells **(Fig. 2A)**, which appears to correlate with the overall cellular levels of VPS4B. However, the percentage of EEA1 positive puncta positive for VPS4A upon EGF treatment was not much different between control and CPAP-depleted cells, even with the diminished cellular levels of this protein in CPAP depleted cells **(Fig. 2B)**. Of note, the degree of localization of VPS4A onto EE upon EGF treatment is limited compared to the degree of VPS4B recruitment (Fig. S4), suggesting that the EE localization dynamics of VPS4A may be different from that of VPS4B. Since our studies suggest that CPAP is associated with the EE localization of only VPS4B even when the cellular levels of both VPS4A and VPS4B are affected, rest of our studies were focused on the impact of CPAP on VPS4B.

**Fig. 2.**
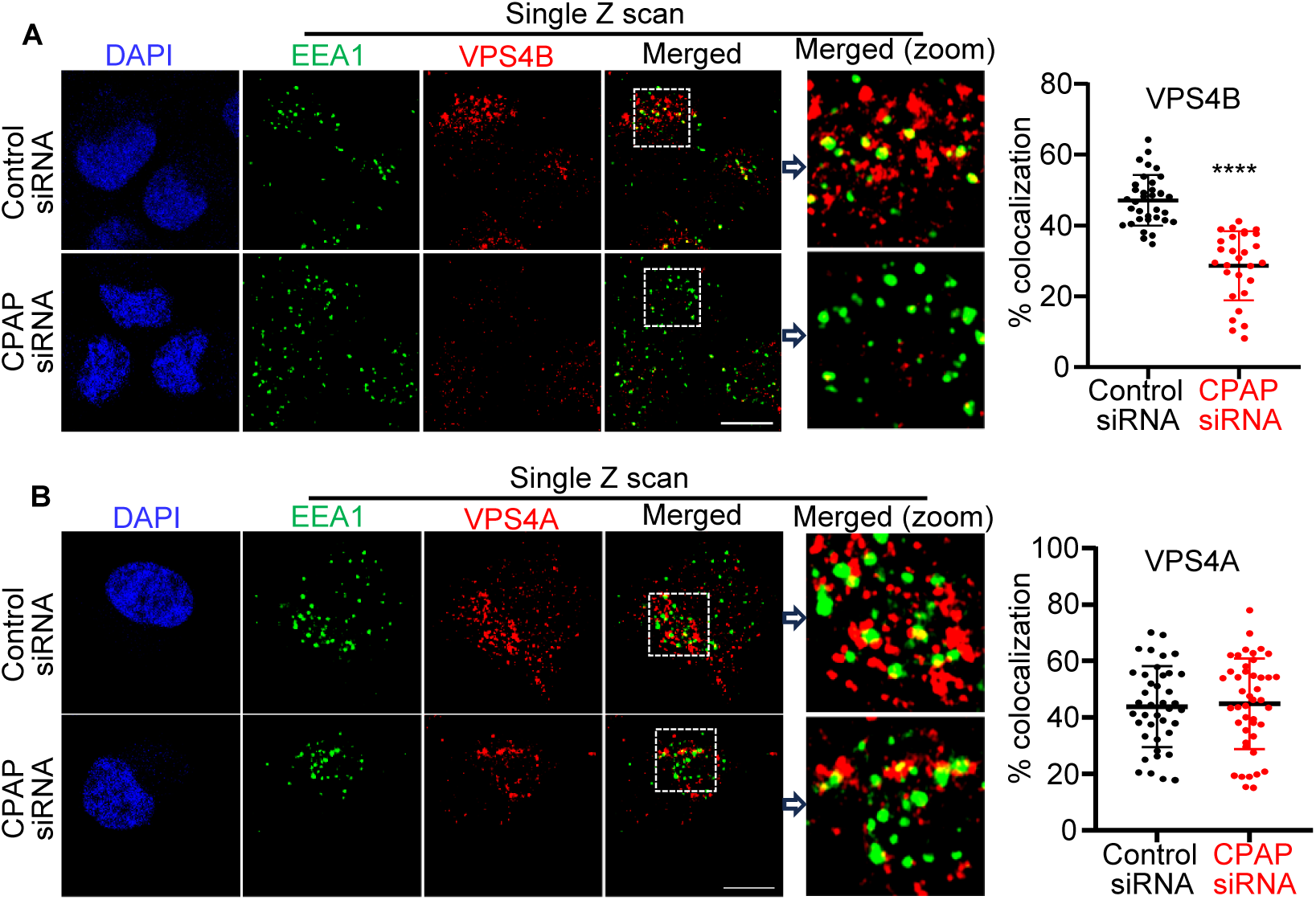
Endosome localization of VPS4B is disrupted in CPAP depleted cells. HeLa cells expressing control or pooled CPAP-siRNA were treated with untagged EGF for 60 min, stained for ESCRT pathway protein paralogs, VPS4B **(A)** and VPS4A **(B)** along with EEA1 (to mark EE), and imaged by Airyscan super-resolution microscopy. Left panels: representative single Z-plane of images showing localization of VPS4B or VPS4A on EEA1-positive puncta. Right panels: colocalization (yellow) was quantified by counting percentages of EEA1-positive (green) puncta containing VPS4 protein-positive (red) puncta in representative single Z-planes of each cell and quantified from multiple cells across at least 3 experiments. Object-based colocalization macro tool of FIJI was employed. Scale bar: 10µm. *p*-values by Mann-Whitney test. ****<0.0001.

### Over-/exogenous-expression of CPAP enhances cellular levels of VPS4B

Diminished CPAP expression appears to cause reduction in the protein levels of VPS4B and suggested that cellular levels of VPS4B may be proportionately increased or decreased depending on the CPAP protein levels. Therefore, to examine if CPAP overexpression increases the cellular levels of VPS4B, HEK293T cells were transfected with different amounts of GFP-tagged-CPAP cDNA and subjected to IB. As observed in **Fig. 3A**, increasing GFP-CPAP expressions resulted in a corresponding increase in the cellular levels of VPS4B. Further, exogenous expression of RNAi-resistant GFP-CPAP in CPAP-depleted cells, by doxycycline (doxy) treatment, caused the higher expression of VPS4B in endogenous CPAP-depleted cells as compared to CPAP-depleted control cells **(Fig. 3B)**. These observations, in association with the observations of Fig. 1A, confirm that the cellular level of VPS4B, perhaps its stability, is specific to and dependent on CPAP protein levels.

**Fig. 3.**
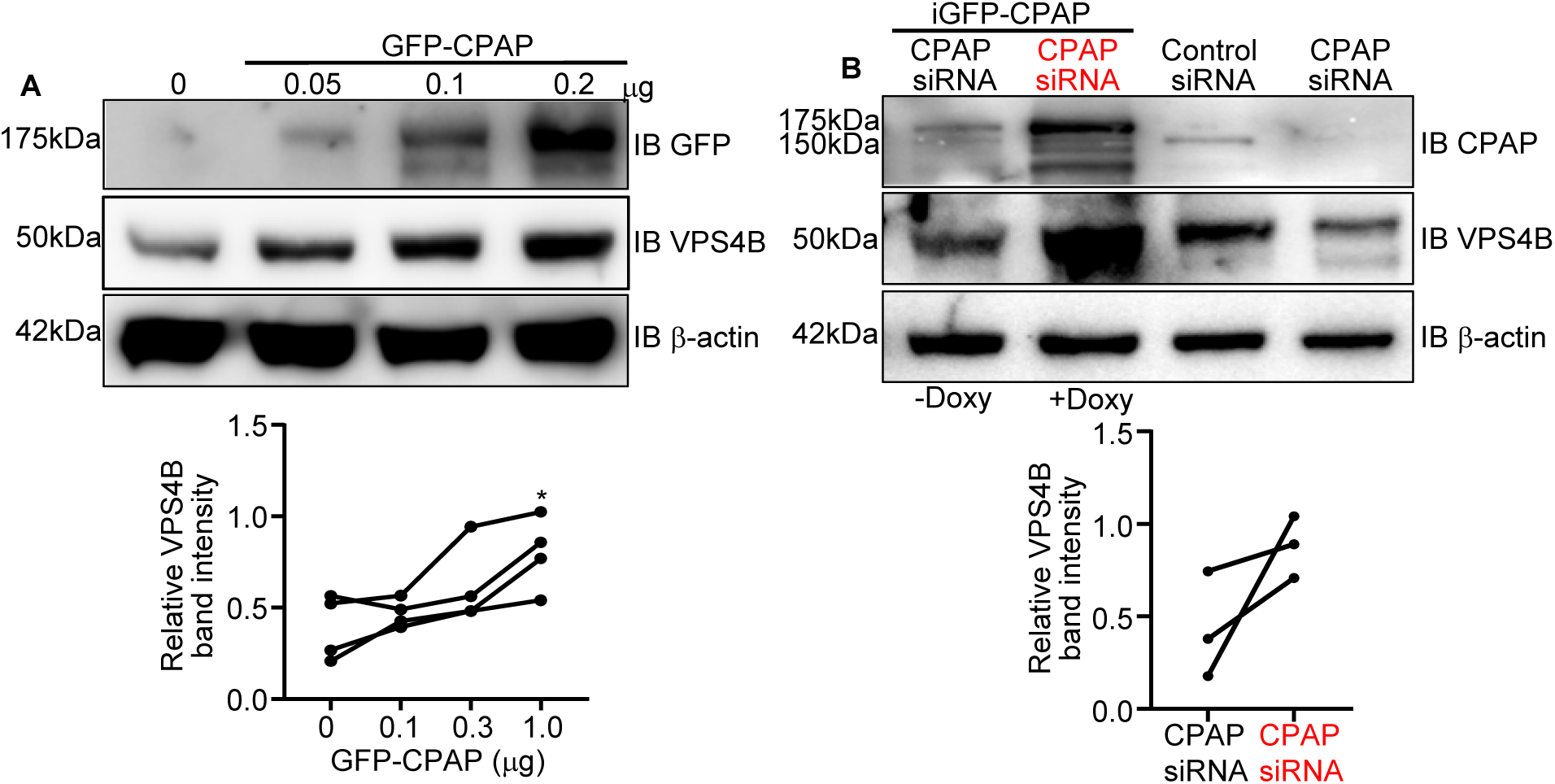
Over-/exogenous-expression of CPAP enhances cellular levels of VPS4B. **(A)** HEK293T cells were transfected with indicated amounts of GFP-CPAP plasmid for 24h and subjected to immunoblotting using anti-GFP and -VPS4B antibodies. Representative blots (upper panel) and densitometric values of VPS4 protein bands (relative to β-actin) from at least three independent experiments (lower panel), are shown. **(B)** Control-siRNA or CPAP-siRNA3 (compatible with RNAi-resistant cDNA vector constructs) treated HEK293T cells, with and without doxy-inducible, RNAi-resistant GFP-CPAP expression, were immunoblotted using CPAP-and VPS4B-specific antibodies. Representative blots (upper panel) and densitometric values of VPS4 protein bands (relative to β-actin) from at least three independent experiments (lower panel), are shown. *p*-values by paired *t*-test. *<0.05.

### Reintroduction of CPAP and overexpression of IST-1 fully restores EE recruitment of VPS4B in CPAP depleted cells

Since CPAP depletion appears to have disrupted residual VPS4B localization to the EE and CPAP overexpression increased the cellular levels of VPS4B, we examined if reintroduction of CPAP expression alone can restore the endosome localization of VPS4B in CPAP-deficient cells. HeLa cells stably expressing doxycycline-inducible, RNAi-resistant, GFP-tagged-CPAP cDNA were treated with CPAP-specific-siRNA to deplete endogenous CPAP expression and doxy to induce exogenous GFP-CPAP expression. These cells were treated with EGF to induce EVT, subjected to immunostaining with anti-EEA1-and -VPS4B antibodies and Airyscan super-resolution microscopy as depicted in **Fig. 4A**. Correlating with the protein levels and confirming the observations of Fig. 2A, CPAP-siRNA treated control cells showed significantly lower percentage of EEA1 puncta with VPS4B positive puncta localization compared to control siRNA-treated cells. However, the proportion of EEA1-positive structures with VPS4B localization was profoundly increased in CPAP-depleted cells upon re-introduction of CPAP protein **(Fig. 4B)**, suggesting that endosomal recruitment of VPS4B is enhanced by restoring CPAP-dependent cellular pool of VPS4B. Of note, our recent report has shown that GFP-CPAP expression in CPAP-siRNA treated cells also restores endosomal trafficking of EGFR cargo (24).

**Fig. 4.**
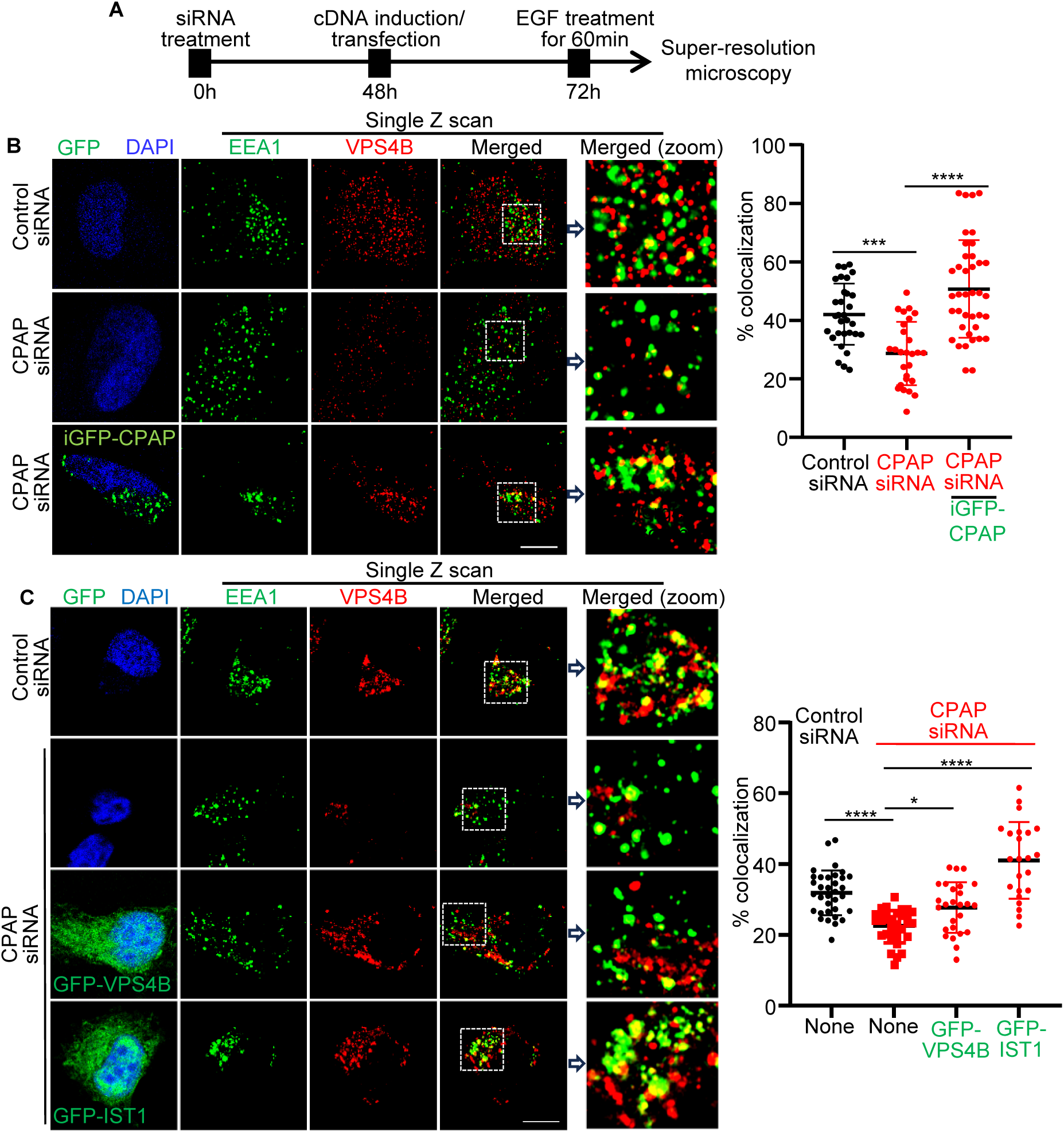
Reintroduction of CPAP and overexpression of IST-1 fully restores EE recruitment of VPS4B in CPAP depleted cells. **(A)** Schematic depicting experimental strategy for panels D and E using HeLa cells. CPAP-siRNA treated cells were transfected with iGFP-CPAP **(B)** and GFP-VPS4B or GFP-IST1 **(C)** and the cells were stained for EEA1 (using AF-647-linked secondary antibody) and VPS4B (using AF-568-linked secondary antibody) and imaged by Airyscan super resolution microscopy. Only GFP-positive cells were imaged for GFP-tag vector-construct transfected cells. D&E left panels: representative single Z-plane of images showing localization of VPS4B on EEA1-positive puncta. D&E right panels: colocalization (yellow) was quantified by counting percentage of EEA1-positive (pseudo-color; green) puncta containing VPS4B-positive (red) puncta in representative single Z-planes of each cell and quantified from multiple cells across at least 3 experiments. Object-based colocalization macro tool of FIJI was employed. Scale bar: 10µm. *p*-values by Mann-Whitney test. ****<0.0001.

Since both the cellular level and EE localization of VPS4B are diminished upon CPAP depletion, it is unclear if CPAP is required for recruiting VPS4B on to endosome or if the diminished recruitment is due to its lowered cellular levels alone. We reasoned, if CPAP plays a role only in VPS4B stability but not in its EE localization, that restoration of VPS4B protein levels may restore its recruitment to EE in CPAP-deficient cells. This was tested by overexpressing VPS4B in CPAP-depleted cells. Importantly, previous reports have shown that the ESCRT-III subunit IST1 exists in two different pools on endosomes and regulates VPS4 localization and assembly on endosomes (30–32). Therefore, the effect of IST1 overexpression on endosomal localization of VPS4B in CPAP-depleted cells was also tested. As observed in **Fig. 4C**, a partial restoration of the localization of VPS4B on EE in CPAP-depleted cells, as indicated by the percentage of EEA1 positive structure with VPS4B, was observed upon exogenous/over-expression of VPS4B suggesting that CPAP may have a role in recruiting VPS4B on to endosome. Interestingly, overexpression of IST1 resulted in not only an increase in the amount of VPS4B puncta, but also a robust reversal of CPAP depletion-associated disruption of VPS4B recruitment onto EE. This suggests that higher cellular levels of IST1 can enhance the EE localization of VPS4B in CPAP-depleted cells. Importantly, while both VPS4B and IST-1 overexpression caused a similar increase in the abundance of VPS4B-positive puncta in CPAP-depleted cells, the percentage of VPS4B-positive EE structures were profoundly higher in IST-I overexpressing cells compared to VPS4B overexpressing cells. This suggests that higher levels of IST-I can compensate for CPAP deficiency in terms of VPS4B localization on to endosomes. Of note, our observations that CPAP depletion has no effect on IST1 cellular levels **(Fig. S5A)** and IST1 depletion diminishes only the EE localization, but not the cellular levels, of VPS4B **(Fig. S5B)** suggest that CPAP and IST1 may be facilitating EE localization of different pools of VPS4B.

### CPAP depletion disrupts cellular processes such as exosome release, viral budding, and cytokinesis

Our results suggest that CPAP is involved in maintaining the cellular pool of VPS4B for facilitating MVB biogenesis, independently of its upstream ESCRT-0 function, which involves the localization of TSG101 to EE (24). Importantly, AAA+ ATPase function of VPS4 proteins is critical for a multitude of cellular processes including exosome release, viral budding and cytokinesis (33–36). Therefore, we hypothesized that loss of CPAP, which causes VPS4B loss, could negatively impact the cellular processes that are dependent on VPS4B. To test this notion, control and CPAP-deficient and CPAP-overexpressing cells were examined for their ability to produce extracellular vesicles. **Fig. 5A** shows that CPAP-depleted cells produced significantly lower amounts of extracellular vesicles compared to control cells. Reciprocally, consistent with the higher protein levels of VPS4B upon CPAP-overexpression (Fig. 3A), the overall amounts of extracellular vesicles released by GFP-CPAP-expressing HEK293T cells were higher than that by control cells. We also tested the ability of CPAP-depleted cells to generate (produce and release) HIV-based lentivirus particles, which is known to be VPS4B-dependent (16). Electron microscopy images of lentiviral expression-and packaging vector-transfected cells suggested not only a difference in the size of virus-like particles between control and CPAP-depleted cells, but also there appears to be a defect in viral egress, as indicated by large undetached structures and fewer number of free virus-like particles, in CPAP-depleted cells **(Fig. 5B)**. The ability of control and CPAP-depleted cells to produce comparable amounts of lentiviral components, as indicated by IB to detect the HIV GAG protein (p55), suggests that the amounts of viral protein produced by control and CPAP-depleted cells are comparable. To quantify the released functional virus-like particles, fresh cells were treated with lentivirus-containing supernatant from GFP expression and packaging vector-transfected control and CPAP-depleted cells and examined for GFP expression levels. Cells transduced with lentivirus from CPAP-depleted cells showed profoundly lower GFP expression compared to that of control cells. This suggests that CPAP deficient cells released lesser amounts of functional virus-like particles. CPAP-depleted HeLa cells were also examined for the dynamics of cytokinesis, another VPS4B-dependent process, by assessing the midbody resolution timing by live imaging. **Fig. 5C** shows that cell separation/mid body abscission time during cell division was significantly delayed in CPAP-depleted cells compared to control cells and the average cell separation time more than doubled upon CPAP depletion. Overall, these results, in association with Figs. 1-4, confirm that CPAP deficiency not only causes destabilization of VPS4B protein levels, but also leads to compromised VPS4B-dependent cellular processes.

**Fig. 5.**
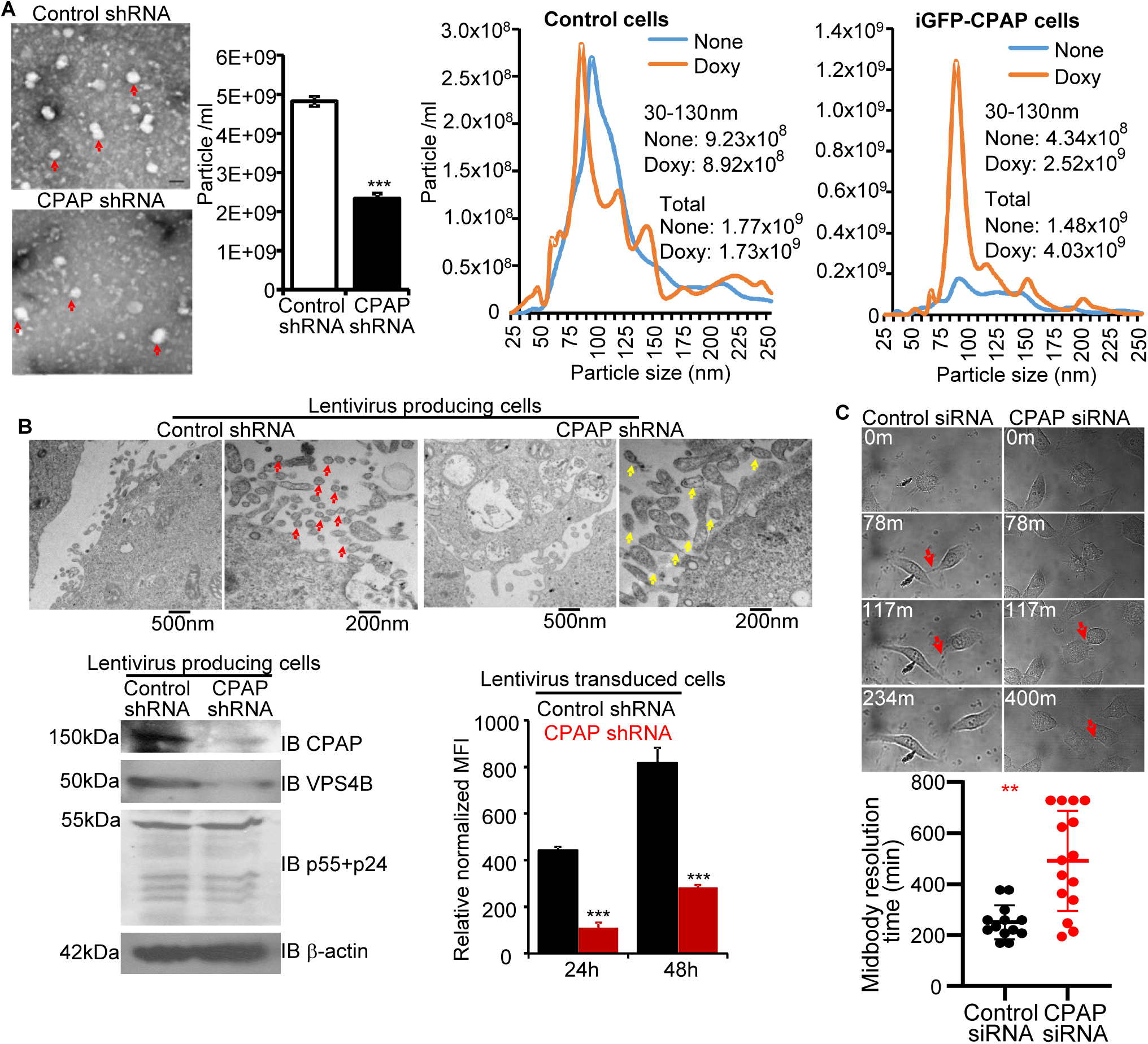
Extracellular vesicle release, viral budding and cytokinesis are influenced by CPAP levels. **(A)** HEK293T cells stably expressing control or CPAP -shRNA lentiviral vectors were rinsed with serum-free medium and equal number of cells were cultured in DMEM media containing exosome-depleted FBS for 48h. Culture supernatants were collected and clarified by centrifugation at 4,500 RPM followed by 0.22μm filtration. Left panel: the supernatants were subjected to PEG precipitation and reconstituted in proportionate volume of PBS, adsorbed on to carbon formvar-coated copper grids, negatively stained with 1% uranyl acetate, and imaged by transmission electron microscopy (TEM). Representative images with examples of exosome-sized particles indicated by red arrows are shown. Middle panel: clarified and filtered (unconcentrated), culture supernatants were subjected to nanoparticle tracking assay (NTA) using NanoSight instrumentation and the abundance of particles with 30-130nm size-range (mean±SD of values from 3 parallel cultures) are shown. Right panel: Control and doxy-inducible GFP-CPAP (iGFP-CPAP) -expressing HEK293T cells were cultured with and without doxy as described above, and the clarified supernatants were subjected to NTA. Histogram overlays of a representative assay that was carried out in triplicate with details on the abundance of particle sizes between 30-130nm and 25-250nm (total) are shown. **(B)** HEK293T cells stably expressing control or CPAP-shRNA were transfected with GFP-encoding lentiviral vector pLL3.7 along with packaging vectors (pMD2.G, pRSV-Rev and pMDLg/pRRE) as described under Material and Methods to generate lentiviral particles, and the virus containing supernatants and cells were harvested after 24h. Upper panel: representative TEM images of transfected lentivirus-producing cells showing large particles/vesicular structures undetached in CPAP-depleted cells (yellow arrows) as compared to the free smaller virion-like particles (red arrows) close to control cells. Lower left panel: Lysates of transfected cells were subjected to WB to detect viral GAG protein p55 (antibody against p55+p24; One World lab) along with CPAP and VPS4B and IB images are shown. Lower right panel: HEK293T cells were transduced with cell-free supernatants from 3 parallel cultures, at 1:4 dilution, and cultured for 48h, analyzed by FACS to determine GFP expression, and the mean±SD of mean fluorescence intensity (MFI) values of these transduced cells, normalized by dividing with transfection efficiency values (GFP-positive cell percentage of virus-producing cells), are shown. **(C)** HeLa cells were treated with control and pooled CPAP -siRNA for 48h and subjected to live cell imaging to detect cytokinesis. Representative cells undergoing division/midbody (indicated by red arrow) separation (upper panel) and quantification of midbody resolution time of multiple cells (lower panel) are shown. *p*-values by Mann-Whitney test. **p<0.01; ***p<0.001.

### Functional domain(s) of CPAP that promote stability and EE localization of VPS4B are located in its C-terminus

To begin to understand if specific functional domain(s) of CPAP are responsible for the stability and EE localization of VPS4B, RNAi-resistant CPAP cDNA constructs with major, previously described, functional domain deletions **(Fig. S6)** were expressed in HeLa cells and studied for VPS4B protein dynamics. GFP-tagged, doxy-inducible, RNAi-resistant CPAP-FL (full-length CPAP), CPAP-ΔPN2-3 (CPAP with tubulin-binding domain deletion), CPAP-ΔCC4 (CPAP with coiled-coiled region 4/centriole localization domain deletion), and CPAP-ΔTCP (CPAP with STIL binding-and microcephaly-mutation associated domain deletion) constructs were described before (26). Cells that express these cDNA vector constructs were subjected to transient endogenous CPAP depletion using RNAi approach, treated with doxy to induce exogenous proteins and examined for VPS4B levels. As observed in **Fig. 6A**, CPAP-FL expression, under endogenous CPAP depletion, increased the cellular levels of VPS4B when compared to control cells suggesting that exogenously expressed CPAP causes stabilization of cellular VPS4B levels as observed in Fig. 3. While CPAP-ΔPN2-3 and CPAP-ΔTCP mutant constructs showed similar stabilizing effects to that of CPAP-FL on VPS4B, the exogenous expression of CPAP-ΔCC4 failed to cause accumulation of VPS4B levels, suggesting that the CC4 domain of CPAP is required for stabilizing VPS4B levels.

**Fig. 6.**
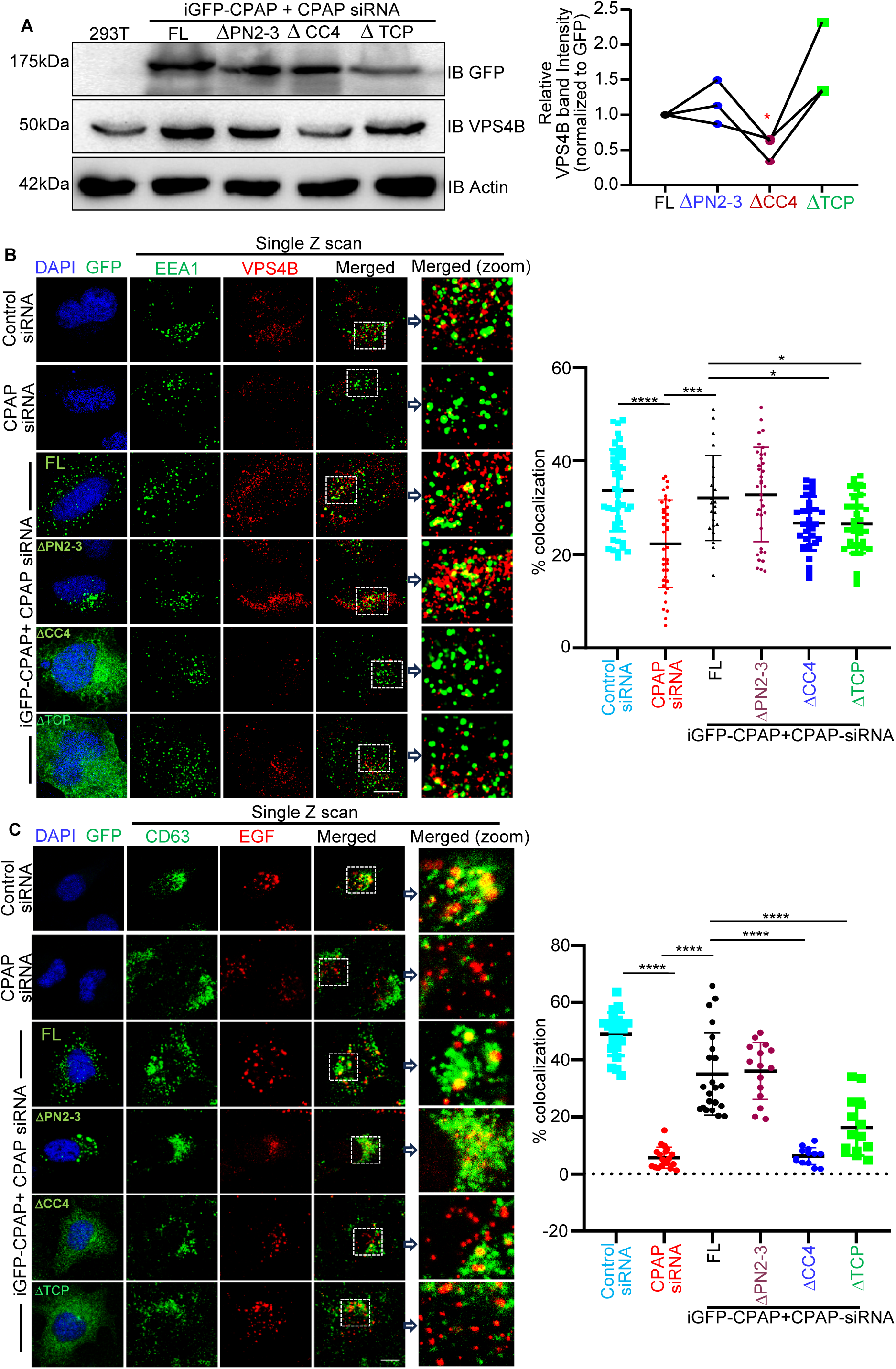
Functional domains of CPAP that promote stability and EE localization of VPS4B are located at its C-terminus. **(A)** HEK293T cells stably expressing doxy-inducible, RNAi resistant, FL, ΔPN2-3, ΔCC4 or ΔTCP versions of GFP-CPAP were treated with CPAP-siRNA3 for 24h, followed by treatment using doxy for another 24h. Representative IB of cell lysates (left panel), and relative densitometric values of VPS4B bands from three independent experiments with FL group value as base value of 1 for each experiment (right panel) are shown. **(B)** HeLa cells expressing Control-siRNA or CPAP-siRNA3, and CPAP siRNA-resistant GFP-CPAP-FL, ΔPN2-3, ΔCC4 or ΔTCP expression vectors were treated with untagged EGF ligand for 60m, stained for EEA1 (using AF-647 linked secondary antibody) and VPS4B (using AF-568 linked secondary antibody) and imaged using Airyscan super-resolution microscopy. Only GFP-positive cells were imaged for GFP-tagged construct transfected cells. Left panel: representative single Z-plane of images showing localization of EEA-1 positive puncta with VPS4B-positive puncta. Right panel: colocalization (yellow) was quantified by counting percentages of EEA1-positive (pseudo-color; green) puncta containing VPS4B-positive (red) puncta in representative single Z-planes of each cell and quantified from multiple cells across at least 3 experiments. **(C)** HeLa cells expressing control-siRNA or CPAP-siRNA3, and CPAP-siRNA-resistant GFP-CPAP-FL, ΔPN2-3, ΔCC4 or ΔTCP expression vectors were treated with AF555-EGF for 60 min and stained for CD63 (using AF-647 linked secondary antibody) to mark MVBs/late endosomes and imaged by Zeiss 880 confocal microscope to determine AF555-EGF and CD63 colocalization. Only GFP positive cells were imaged for GFP-tagged protein expressing cells. Left panel: representative single Z-plane of images showing localization of CD63-and EGF-positive puncta. Right panel: colocalization (yellow) was quantified by counting percentages of EGF-positive (red) puncta containing CD63-positive (pseudo-color; green) puncta in representative single Z-planes of each cell and quantified from multiple cells across at least 3 experiments. Object-based colocalization macro tool of FIJI was employed. Scale bar: 10µm. *p*-value by paired t-test for panel A and Mann-Whitney test for panels B&C. *<0.05, ***<0.001, ****<0.0001.

Next, we examined the impact of expressing mutant cDNA constructs of CPAP on VPS4B localization to EE. As shown in **Fig. 6B**, induction of exogenous expression of CPAP-FL and CPAP-ΔPN2-3, but not CPAP-ΔCC4 and CPAP-ΔTCP, in endogenous CPAP-depleted cells restored EE localization of VPS4B. While Fig. 6A shows that only the CC4 domain is required for stabilizing the cellular levels of VPS4B, this observation suggests that C-terminal domains of CPAP demonstrate a differential role, one for protein stability and the other for facilitating the EE localization of VPS4B. We have then tested the impact of exogenous expressions of full-length and mutant CPAP proteins, in CPAP-depleted cells, on the downstream event, i.e. transport of internalized receptor cargo to late endosomes (LE) by examining EGF-AF555 ligand-treated cells for the degree of EGF localization to the CD63-positive (MVB/LE) vesicular structures. As observed in F**ig. 6C**, CPAP-FL-and CPAP-ΔPN2-3-expressing cells, with endogenous CPAP depletion, show higher localization of EGF-AF555 to CD63-positive structures compared to CPAP-depleted cells indicating the restoration of endosomal cargo transport. On the other hand, the degree of EGF localization on CD63-positive structures in CPAP-ΔCC4-or CPAP-ΔTCP-expressing cells was not considerably different from that of CPAP-depleted cells. This suggests that endocytic trafficking function is not restored in CPAP-depleted cells in the absence of C-terminal domains CC4 and TCP in the exogenously expressed protein, potentially due to diminished cellular levels and EE localization/function of VPS4B, respectively.

### VPS4B interacts with CPAP and is targeted for proteasome degradation under CPAP depletion

To further understand how CPAP contributes to the stability and endosome localization of VPS4B, we tested if this AAA+ ATPase forms a protein complex with CPAP. HEK293T cell lysates were subjected to immunoprecipitation (IP) using VPS4B-specific antibody followed by IB detection of VPS4B and CPAP. As shown in **Fig. 7A**, CPAP was detected in IP of VPS4B suggesting that these proteins are indeed present in a physiological complex. To substantiate this observation of a protein-protein interaction between CPAP and VPS4B, lysates of GFP-CPAP-expressing cells were also subjected to IP and IB. **Fig. 7B** shows that GFP (CPAP)-specific IP revealed VPS4B along with GFP-CPAP in them, confirming an interaction/complex formation between these two proteins.

**Fig. 7.**
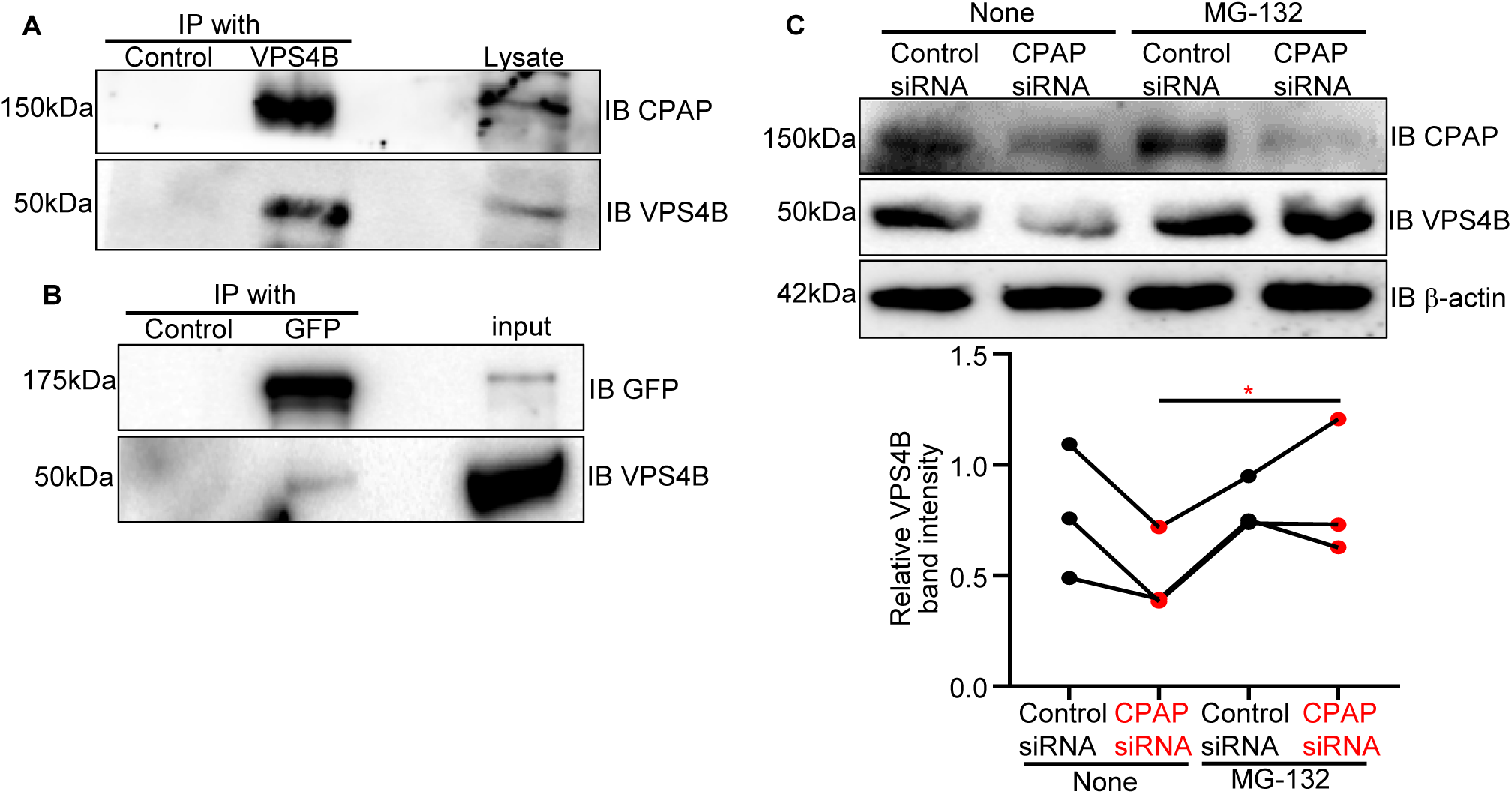
VPS4B interacts with CPAP and is targeted for proteasome degradation under CPAP depletion. **(A)** Lysates of HeK293T cells were subjected to IP using control or anti -VPS4B antibody beads, followed by IB using indicated antibodies. Representative blots of three independent experiments are shown. **(B)** Doxy-inducible GFP-CPAP expressing HEK293T cells treated with doxy for 24h were subjected to IP using control or anti-GFP antibody beads, followed by IB using anti-GFP and -VPS4B antibodies. Representative blots of three independent experiments are shown. (**C)** HEK293T cells treated with control-or CPAP-siRNA for 48h and MG-132 (or DMSO containing medium control; none) for another 24h and subjected to IB with anti-GFP and -VPS4B antibodies. Representative blots (upper panel) and densitometric values of VPS4 protein bands (relative to β-actin) from at least three independent experiments (lower panel), are shown. *p*-value by paired t-test *<0.05.

Diminished cellular levels of VPS4B in CPAP-depleted cells indicate that VPS4B is degraded in the absence of interaction with CPAP. Therefore, we examined if VPS4B protein is targeted for proteasome degradation in the absence of CPAP. Control and CPAP-depleted cells were treated with proteasome inhibitor, MG-132 and the cell lysates were subjected to IB. As observed in **Fig. 7C**, while MG132-treatment did not have much impact on the VPS4 levels of control cells, proteasome inhibition resulted in accumulation of VPS4B protein in CPAP-depleted cells. This suggests that complex formation with CPAP prevents proteasomal degradation of VPS4B. Overall, our observations suggest that CPAP deficiency results in targeting the cytoplasmic pool of VPS4B for proteasomal degradation and its diminished localization/function on EE.

### Cellular stability of VPS4B is dependent on its interaction with the C-terminal CC4 domain of CPAP

Since the C-terminal CC4 domain appears to have the VPS4B-stabilizing property (Fig. 6A), we examined if previously described CPAP fragments (37) and functional domain mutants (26), depicted in **Fig. S6**, interact with VPS4B. IP assays using lysates of cells expressing fragments of CPAP showed that C-terminal fragments (CP-3), but not N-terminal (CP-1) or middle fragments (CP-2), pulled down VPS4B protein (**Fig. 8A**) suggesting that the VPS4B interaction motif resides with in the C-terminus. Further, immunoprecipitation of GFP-tagged CPAP-FL and its functional domain-mutant CPAP proteins revealed that CPAP-ΔCC4, but not CPAP-ΔPN2-3 or CPAP-ΔTCP, mutant protein lacked the ability to pull down VPS4B (**Fig. 8B**). Since CPAP-ΔCC4 mutant expressing cells with endogenous CPAP depletion showed lower VPS4B levels (Fig. 6A), these cells were also subjected to MG-132 treatment, to inhibit proteasome-mediated protein degradation, and examined for VPS4B levels. While the induction of exogenous CPAP-FL protein under endogenous CPAP depletion resulted in an increase in VPS4B protein levels, induction of ΔCC4 mutant expression did not cause an increase in VPS4B protein level **(Fig. 8C)**. On the other hand, MG-132 treatment caused the accumulation of VPS4B in CPAP-ΔCC4 expressing cells suggesting that lack of CC4 domain function of CPAP causes constitutive proteasome targeting of VPS4B for degradation. Overall, these observations reiterate that cellular levels of VPS4B are stabilized/maintained by interaction with the CC4 domain of CPAP, which prevents its targeting for degradation by the proteasome.

**Fig. 8.**
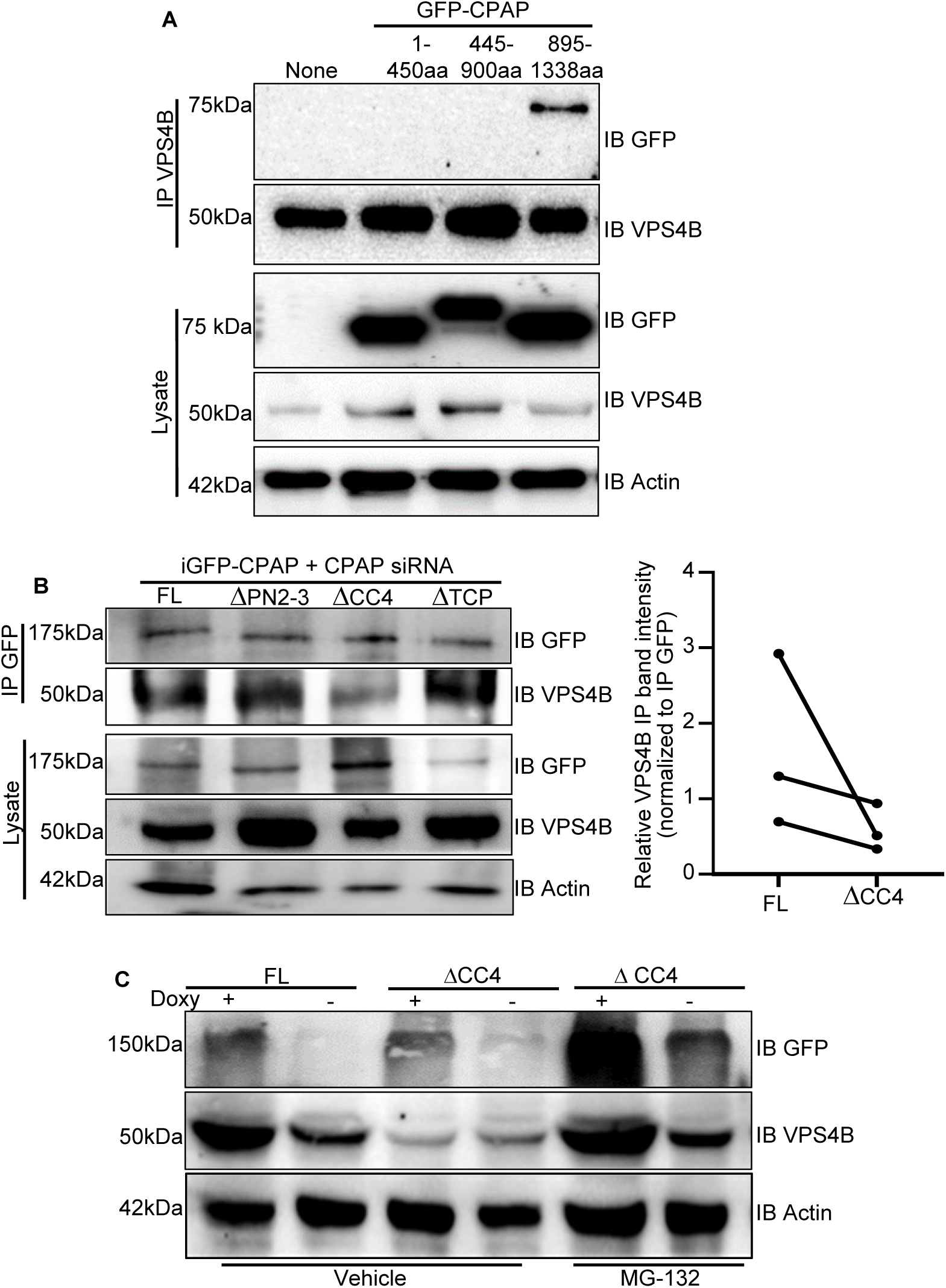
Cellular stability of VPS4B is dependent on its interaction with the C-terminal CC4 domain of CPAP. **(A**) HEK293T cells that were transfected with expression vectors encoding N-terminal (1-445 aa), middle (445-900 aa) or C-terminal (895-1338 aa) fragments of CPAP for 24h and subjected to IP using VPS4B antibody and immunoblotted using anti-GFP and -VPS4B antibodies. Representative blots of 3 independent experiments are shown. **(B)** HEK293T cells stably transfected with doxy-inducible RNAi resistant FL or ΔPN2-3 or ΔCC4 or ΔTCP versions of GFP-CPAP plasmids were treated with CPAP siRNA3 for 24h, followed by doxy for another 24h, and subjected to IP using anti-GFP antibody and IB using anti-GFP and -VPS4B antibodies. Representative blots (left panel) and densitometric values of VPS4 protein bands (relative to β-actin; FL vs ΔCC4 group comparison) from at least three independent experiments (right panel), are shown. **(C)** HEK293T cells expressing RNAi-resistant GFP-CPAP-FL or -ΔCC4 were treated with CPAP-siRNA3 for 48h to deplete endogenous protein and doxy for another 24h to express exogenous protein. MG-132 (or DMSO containing medium control; none) were added to the cultures along with doxy. Cell lysates were subjected to IB using anti-GFP and -VPS4B antibodies and representative images from 3 independent experiments are shown.

## Discussion

Centriole-and microcephaly-associated protein, CPAP (Centrosomal P4.1-associated protein) has a well-documented role in the regulation of centriole duplication (19,20,26). Mutations in the CENPJ gene in human patients have been associated with microcephaly and Seckel syndrome, and disruption of mouse *Cenpj* gene expressed phenocopies Seckel syndrome (38–41). CPAP is an α-tubulin-binding protein and this tubulin-binding property (42–44) is important for its function on the centrioles, especially in restricting the centriole length to <500nm (18–20,44) as well as mitotic spindle orientation during cell divisions (26). Centrioles serve as basal bodies, which template the formation of cilia (45). CPAP has also been shown to regulate the length of cilia (18,20,21). We have uncovered a non-canonical function of CPAP as a positive regulator of endosomal transport and lysosomal degradation of endocytosed cell surface receptor EGFR that limits EGFR function and prevents EMT of cancer cells (23,25). Further, our recent report showed that CPAP plays an essential role, similar to the ESCRT-0 protein HRS, in recruiting TSG101 (ESCRT-I protein) onto EE and facilitating the downstream events during MVB formation and EGFR cargo sorting (24), providing the mechanistic explanation for our previous report (23) on its role as a positive regulator of endosomal transport. Importantly, endocytosis of lysosome-targeted cell surface receptors like EGFR leads to the sequential recruitment of other ESCRT protein complexes (ESCRT-I and ESCRT-III) and accessory proteins, including ALIX, VPS4A and VPS4B, to the EE. These downstream ESCRT proteins, particularly ESCRT-III members and AAA+ ATPase VPS4, have also been implicated in MVB formation (15). VPS4 function is critical for the dissociation and reassembly of ESCRT-III subunits on endosomal membrane surface during ILV/MVB formation (29). ESCRT-III filaments constrict and cut the membrane neck to form ILVs (13) and VPS4 proteins promote the dissociation/reassociation of ESCRT-III complex until membrane scission is achieved (29). However, much remains to be learned about the endosomal recruitment and functional dynamics of ESCRT-III complex and VPS4 proteins.

Here, we report that CPAP is critical for the protein stability, endosome localization, and MVB formation function of VPS4B, and this role of CPAP is independent from its role as an ESCRT-0 in recruiting ESCRT-I protein TSG101. While we found that the cellular levels of both VPS4 proteins are affected by CPAP levels, EE recruitment of residual VPS4B paralog, but not VPS4A, is diminished in CPAP-depleted cells. Further, we found that the degree of localization of VPS4A onto EE upon EGF treatment is limited compared to the degree of VPS4B recruitment. This suggests that the EE localization dynamics of VPS4A may be different from VPS4B and VPS4B may have a dominant role in MVB formation than VPS4A. In this regard, while both paralogs of human VPS4 are thought to be involved in intracellular protein trafficking (46), they are differentially expressed in various organs suggesting that they could have distinct functional properties (47). Further, co-expression of VPS4A and VPS4B proteins revealed only partial co-localization (47). Overlap and differences in the role of VPS4A and VPS4B in cytokinesis, an ESCRT-pathway dependent event, was also recently reported (48). Based on the differential expression profiles of these two proteins in various tumors and loss and gain-of-function studies, it has been suggested that the regulation of gene expression and protein stability of VPS4A and VPS4B occur independently (49). Our observations here support the notion that these protein paralogs have distinct functional roles in endosomal trafficking of cargo proteins.

Although both the cellular level and EE localization of VPS4B are diminished upon CPAP depletion, the role of CPAP in recruiting VPS4B onto EE required further validation because the diminished recruitment can merely be due to its lowered cellular levels in CPAP depleted cells. Our observation that VPS4B overexpression in CPAP-depleted cells could restore its EE localization only partially suggests that CPAP does facilitate VPS4B recruitment on to EE. Importantly, previous reports have shown that the ESCRT-III subunit IST1 exists in two different pools on endosomes and it facilitates VPS4 localization and assembly on endosomes (30–32). Our observations that CPAP depletion has no effect on IST1 cellular levels, IST1 depletion diminishes only the EE localization, but not the cellular levels, of VPS4B, and IST1 overexpression in CPAP-depleted cells causes a robust increase in VPS4B recruitment to endosome suggest that CPAP and IST1 may be facilitating EE localization of different pools of VPS4B. Overall, while different modes of regulation of VPS4 localization, assembly and function by IST1 have been known (30,31,50), our findings suggest that CPAP is not only essential for VPS4B stability, but may also be involved in EE localization/maintenance of VPS4B protein.

Our observations show that CPAP is involved in maintaining the cellular pool of VPS4B for facilitating MVB biogenesis, independently of its upstream ESCRT-0 function, which involves localization of TSG101 to EE (24). Previous reports have shown that diminished VPS4B causes defective cargo transport, leading to persistent signaling by internalized EGFR (51). Our reports have shown that CPAP deficiency causes similar defective late endosome and lysosome targeting of degradative cargo and persistent signaling by internalized EGFR leading to epithelial mesenchymal transition and enhanced tumorigenicity of cancer cell lines (23,25). Hence, as the AAA+ ATPase function of VPS4 proteins is critical for a multitude of cellular processes including exosome release, viral budding and cytokinesis (33–36), we hypothesized that CPAP deficiency will disrupt these cellular processes due to the loss of VPS4B protein. In fact, although it is yet to be determined if the effects are VPS4B loss-associated by restoring its cellular levels, CPAP deficiency caused the suppression of extracellular vesicle release, retrovirus egress and a delay in midbody resolution during cytokinesis. This shows that CPAP deficiency, which causes VPS4B protein loss, can negatively affect all VPS4B-dependent cellular functions irrespective of the ESCRT dependency. Our previous reports on the defective MVB formation and lysosomal targeting of EGFR and EE recruitment of TSG101 to EE (23,24) and the current observations of diminished cellular levels and defective EE recruitment of VPS4B along with defective other VPS4B-dependent cellular processes in CPAP-depleted cells corroborate the notion that this protein can independently regulate different components of the ESCRT machinery.

Other than a previous study which has shown that VPS4B can undergo ubiquitination and proteasomal degradation causing diminished EGFR degradation under hypoxic condition (51), how cellular levels of VPS4B are regulated is unknown. Our observations suggest that, in the absence of direct physical interaction with CPAP, VPS4B undergoes proteasomal degradation. This suggests that the cytoplasmic pool of VPS4B, at least an independent pool, exists in association with CPAP and its functional site recruitment may not be fully dependent on CPAP as suggested by the robust increase in EE localization of VPS4B upon IST1 overexpression in CPAP-depleted cells. As indicated by the increase in VPS4B puncta abundance in CPAP-depleted cells upon IST1 expression, it is also possible that cytoplasmic, but not the functional site stability of VPS4B, may be dependent on CPAP.

Importantly, our observations that VPS4B interacts with, and is recruited to EE by, the C-terminal CC4 and TCP domains of CPAP respectively are suggestive of the additional functional and clinical implications of CPAP-VPS4B interaction beyond MVB formation and other cellular processes described above. For example, CC4 domain of CPAP is also essential for localization of CPAP to its canonical function site on the centrioles (26). On the other hand, the C-terminal TCP domain deletion and mutations are implicated in human microcephaly (26,27). Both TCP and CC4 domain deficiencies can lead to disrupted centriole elongation and centriole duplication (26,28) potentially contributing to ciliopathies and microcephaly. However, whether VPS4B stability and recruitment to endosome are compromised in these clinical conditions in humans are unknown. Nevertheless, our observation that VPS4 protein levels are diminished in cells derived from the CPAP-hypomorphic mice adds a new potential mechanistic dimension to the known clinical phenotypes of CPAP deficiency. Hence, additional studies are needed to determine an association, if any, of VPS4B and ESCRT functions with microcephaly and ciliopathies and for understanding the molecular causes of these CPAP-deficiency/mutation-associated diseases.

Overall, many important cellular membrane fission reactions driven by the ESCRT pathway culminate in the disassembly of ESCRT-III polymers by the AAA+ ATPase VPS4. Here, we demonstrate how a centriole biogenesis protein ensures AAA+ ATPase function, which is critical for MVB formation and endosomal transport of endocytosed cell surface receptor cargo. The newly identified role of CPAP in promoting stability and endosome function of VPS4B appears to be independent from its function as an ESCRT-0, which we described recently (24). Considering the known centriole localization function of CC4 domain and the PCM tethering and STIL binding properties of TCP domain of CPAP (26,28,52), whether the CPAP-VPS4B interaction influences the centriole-and microtubule associated-canonical functions of CPAP remains to be investigated. Additional studies are needed to further identify and dissect the minimum functional domains of CPAP that are involved in regulating VPS4B stability and function and delineate the complex physical and functional interactions between CPAP and other ESCRT pathway proteins. Our finding that CPAP, a critical centriole biogenesis protein, is essential for regulating multiple stages of ESCRT pathway function, could also lead to new research that investigates the complex interactions between ESCRT pathway and variety of cellular events including centriole biogenesis.

### Experimental procedures

#### Cell lines

HEK293T (National Gene Vector Biorepository) and HeLa (ATCC) were used in this study. These cells were cultured in DMEM media supplemented with 10% FBS, sodium pyruvate, sodium bicarbonate, minimum essential amino acids and antibiotics. Transfection of plasmids was performed using calcium phosphate reagent or TransIT X2 reagent from Mirus Bio LLC while siRNA was transfected using the TransIT siquest reagent from Mirus Bio LLC. Exosome-free FBS was purchased from Invitrogen.

#### CPAP-hypomorphic mouse cells

The mouse model with hypomorphic allele of CPAP (*Cenpj^tm/tm^)* was described before (41). This mouse model in C57BL6 background was obtained from The European Mouse Mutant Archive (EMMA) and maintained in our breeding colony. All animal studies were approved by the institutional animal care and use committee of the Medical University of South Carolina. Spleen cells and primary tongue fibroblasts prepared (53) from 8-week-old *Cenpj^+/tm^* (heterozygous) and *Cenpj^tm/tm^* (homozygous; hypomorphic) mice were used in this study.

#### Plasmids and reagents

siRNA/shRNA region targets in CPAP have been described previously (23–25) and these regions were targeted with minor modifications. Briefly, siRNAs targeting 4 different regions of CPAP [CPAP-siRNA1: 5′-CCCAAUGGAACUCGAAAGGAA-3′ (54), CPAP-siRNA2: 5′-UCUAUAUCAUGGCCGACAA-3′ (54), CPAP-siRNA3: 5′-AGAAUUAGCUCGAAUAGAA-3′ (26), and CPAP-siRNA4: 5′-GCUAGAUUUACUAAUGCCA-3′ (23,25)] or scrambled-siRNA control were custom synthesized (by Dharmacon). While individual siRNAs were tested for CPAP knockdown efficiency and off target effects, a pool of these four siRNAs (pooled CPAP-siRNA) were used at a final concentration of 10nM for all experiments where exogenous expression of CPAP protein was not involved. In experiments where exogenous RNAi-resistant CPAP vector constructs are expressed under endogenous CPAP depletion, a single siRNA (CPAP-siRNA3) was used at a final concentration of 10mM. For stable depletion of CPAP, validated lentiviral constructs (pLKO.1) expressing Mission shRNA targeting the region 5′-GCTAGATTTACTAATGCCA-3′ in CPAP or scrambled shRNA (Millipore-Sigma) and used as described in our previous report(23). GFP-CPAP cDNA expression vector and its mutants (CP1-3)(19,28) used in this study were kindly provided by Dr. T.K. Tang, (Academia Sinica, Taipei). RNAi resistant constructs for expressing GFP-CPAP and its domain mutants were kindly provided by Dr. Pierre Gonczy, Swiss Institute for Experimental Cancer Research, Switzerland and the CPAP siRNA targeting sequence has been reported earlier (26). siRNA pools against VPS4A, VPS4B, HRS, and TSG101 were purchased from Santa Cruz Biotech. The vector constructs expressing GFP-VPS4B was kindly provided by Dr. Natalia Elia (National Institute of Biotechnology, Israel) and GFP-IST1 was purchased from Addgene Inc. GFP-encoding lentiviral vector pLL3.7 and the compatible packaging vectors (pMD2.G, pRSV-Rev and pMDLg/pRRE) were purchased from Addgene Inc.

Primary antibodies used in this study: rabbit anti-CPAP (Proteintech; cat#11517-1-AP), mouse anti-CPAP (Abnova, H00055835-M01, for IP studies), anti-GFP (Proteintech; cat#50430-2-AP), anti β-actin (Proteintech; cat# HRP-600008), anti-IST1 (Proteintech; cat# 19842-1-AP), anti-CD63 (BD Biosciences; cat# 556019), anti-EEA1 (Invitrogen; cat#PA1-0337 or BD biosciences; cat#610456), anti-GAG (p55+p24 antibody; One World Labs). Initial studies to evaluate VPS4B levels were performed using antibody provided by Dr. Wesley Sundquist, University of Utah (36). Additional antibodies against VPS4B (cat#sc-377162), HRS (cat#sc-271455), TSG101 (cat#sc-7964) from Santa Cruz Biotech and antibody against VPS4A from Invitrogen (cat#PA5-110573) were also used. Cycloheximide and doxycycline were purchased from Sigma-Aldrich. Unconjugated EGF ligand was purchased from Tonbo biosciences and Alexa fluor-555 conjugated EGF from Invitrogen. Coverslips were mounted using Prolong gold antifade mounting reagent (Invitrogen).

#### Immunofluorescence

Cells grown on coverslips were fixed with 4% paraformaldehyde and permeabilized using 0.1% saponin containing buffer for 30 mins at RT. Blocking with 1% BSA as well as primary and secondary antibody dilutions were made in permeabilization buffer and incubations were done at 37°C. Images were acquired as Z-stacks using either the confocal or super-resolution Airyscan unit of Zeiss 880 microscope using the 63X oil immersion objective with n.a. 1.4 as indicated. Optimal setting as suggested by the software was used to acquire the Z sections.

#### Electron microscopy

HEK293T cells stably expressing control or CPAP shRNA and transfected with GFP-expressing lentiviral vector LL3.7 for 48h, were fixed with 2.5% glutaraldehyde in sodium-cacodylate buffer (Ted Pella Inc) for 30mins and processed as described in (55,56). After dehydration series with alcohol, cells were embedded in Epoxy resin and cured at 60°C for a couple of days. 70-100 nm thin sections on copper grids were examined and imaged using the JEOL 1210 transmission electron microscope (TEM). For exosome analysis, carbon-formvar coated copper grids (Ted Pella, Inc) were floated on drops of exosome solution to adsorb them, followed by 1% uranyl acetate treatment for negative staining.

#### EGFR internalization assay

HeLa cells were rinsed and treated with serum free media for 4h and incubated with media containing cycloheximide [CHX] (5μg/ml) for 1h at 37°C. Chilled cells were treated with CHX media with EGF ligand (50ng/ml) and left for 1h on ice for ligand binding. Cells were washed with chilled serum free media and transferred to 37°C to induce internalization of receptor. For tracking of EGF receptor into vesicles by Zeiss 880 confocal microscopy, Alexa Fluor 555 conjugated EGF (100ng/ml) was used. Untagged EGF was used for other experiments.

#### Immunoblotting (IB) and immunoprecipitation (IP)

Cells harvested at different time-points were lysed using RIPA buffer, supplemented with 0.1M PMSF and protease inhibitor cocktail. Lysates were then spun at 14000 rpm for 30min at 4°C, supernatants were separated and boiled at 95°C for 10m after addition of SDS sample buffer. Post-SDS-PAGE and Western blotting, immunoblotted PVDF membranes were developed using SuperSignal West Dura substrate (Thermo Scientific) chemiluminescence substrate and ChemiDoc imager (Biorad). Anti-rabbit-IgG HRP and -mouse-IgG HRP were obtained from Amersham and Biorad respectively. All blots are representative of at least three independent experiments with similar trends in results. Densitometry analysis was performed using FIJI and presented where appropriate. Immunoprecipitation (IP) with indicated antibodies was subjected to Protein A/G bead (Rockland immunochemicals) pulldown.

#### Lentiviral production and viral transduction

For lentivirus production, HEK293T cells were transfected with lentiviral eGFP encoded vector pLL3.7 and packaging vectors (pMD2.G, pRSV-Rev and pMDLg/pRRE) by calcium phosphate method. Virus-containing media were collected after 24h and 48h, centrifuged at 4500 RPM for 15 min and the supernatants were subjected to 0.22-µm filtration, and serial dilutions were applied on fresh HEK293T cells for 48h. Cells were harvested and analyzed for GFP expression in terms of mean fluorescence intensity (MFI) by flow cytometry. Virus-producing cells were also harvested, 24h post-transfection using pLL3.7 and packaging vectors and subjected to TEM as described above.

#### Extracellular vesicle production assay

HEK293T cells stably expressing control or CPAP -shRNA were generated using lentiviral vectors as described in our previous report (23,25). Cells that are grown to confluency were rinsed with serum-free medium and equal number of cells were cultured in DMEM media containing exosome-depleted FBS for 48h. Culture supernatants were collected and clarified by centrifugation at 4,500 RPM for 30 min followed by 0.22μm filtration. These supernatants were subjected to PEG precipitation on ice for overnight, pelleted by spinning at 4,500 RPM for 30 min. These pellets were reconstituted in proportionate volume of PBS, adsorbed on to carbon formvar-coated copper grids, negatively stained with 1% uranyl acetate, and imaged by TEM as described above. In some assays, clarified and filtered non-concentrated supernatants were subjected to nanoparticle tracking assay (NTA) using NanoSight instrumentation to assess the relative abundances of particles/extracellular vesicles. In some assays, control and doxy-inducible GFP-CPAP-expressing HEK293T cells were cultured with and without doxy as described above for sh-RNA expressing cells, and the supernatants were subjected to NTA.

#### cDNA synthesis and quantitative PCR

Total RNA preparations were made from HEK293T cells that were treated with control or pooled CPAP-siRNA by employing the Trizol method, subjected to cDNA synthesis and used as a template for qPCR using gene specific primer sets. cDNA synthesis was performed using the RevertAid First Strand cDNA Synthesis Kit (Thermo) and qPCR was performed by employing SYBR Green Supermix (Biorad). Relative expression of each factor was calculated first against the house-keeping gene (β-actin; *ACTB*) expression levels, and then the fold change in CPAP-siRNA treated cells as compared to control-siRNA treated cells.

#### Image quantification

Prior to image analysis, Airyscan files were automatically processed using the Zen software (version 2.0) of Zeiss (https://www.zeiss.com/microscopy/us/ products/microscope-software/zen.html). Individual puncta were first delineated, and the background was removed by intensity thresholding in FIJI (https://imagej.nih.gov/ij/). Adjustments were made to entire set of images uniformly within an experiment, before using for quantification of colocalization. Images are presented as maximum intensity projection, or a single Z plane as indicated. Z-stack images were split into single Z planes and, for example: green and yellow pixels were quantified to determine the percentage of colocalization. Object-based colocalization was measured by counting all events of a single Z scan of each cell either manually, or automatically by employing a macro function of FIJI (https://github.com/dsrichardson/fiji_macros/blob/master/2D_object_colocalization), as stated under each figure legend. A single Z-plane of each cell that showed maximum number of puncta was considered for quantification and image presentation. For manual counting, total number of channel 1 puncta (for example green channel) and those showing channel 2 puncta (for example red channel) colocalization (yellow) were counted for the representative single Z-plane of each cell individually using the multi-point tool and counter of FIJI for determining percentage colocalization. The FIJI macro, which is an object-based colocalization tool, identifies the center of mass of each vesicle (with fixed pixels in size for each experiment) in two channels of interest for colocalization analysis of percentage of channel 1 puncta on channel 2 puncta and vice versa. Find Maxima function was used for identifying the center of mass of each vesicle. Uniform maxima values and pixel sizes were used for automatic batch analysis of all images within a single experiment.

#### Statistical consideration

Experiments were repeated several times and all figure panels presented represent at least three experiments that produced similar trends in outcomes. Cumulative values or values from a representative experiment were used for graphical presentation and as indicated in figure legends. Each data point of graphs represents one cell and a single slice from each cell is used for counting all relevant channel puncta. Details on replicated and statistical significance are also mentioned in figure legends. *p*-values were calculated using GraphPad Prism statistical analysis software version 10 (https://www.graphpad.com/scientific-software/prism/). For most comparisons, unpaired non-parametric Mann-Whitney test was used. Densitometric values of IB from multiple experiments were compared by employing paired *t*-test.

## Acknowledgements

We would like to thank Dr. T.K. Tang (Academia Sinica, Taipei) for generously sharing the GFP-CPAP-full length and smaller mutant-expressing cDNA vectors. We extend our gratitude for Dr. Pierre Gonczy (Swiss Federal Institute of Technology, Lausanne, Switzerland) for kindly sharing the doxycycline-inducible vector expressing GFP-CPAP full length and its domain mutants. We would like to thank Dr. Natalia Elia (National Institute of Biotechnology, Israel) for sharing the GPF-VPS4B construct. We would also like to thank Dr. Wesley Sundquist, University of Utah, for sharing several ESCRT pathway related reagents during the early phase of our studies, especially the VPS4B antibody, which set the tone of this paper. This work was supported by National Institutes of Health (NIH) grant R01DE030331 and R21DE026965 to C.V. and R.G. Unrestricted research funds from Hollings Cancer Center and College of Medicine, MUSC supported some aspects of the study. The authors are thankful for the Electron microscopy core and the Cell & Molecular Imaging Shared Resources which is supported by the Hollings Cancer Center, Medical University of South Carolina (P30 CA138313) and the Shared Instrumentation Grant S10 OD018113 as well as the flow cytometry core of MUSC for the FACS instrumentation support.

## Author contributions

R. G. conceived the idea, designed experiments, researched and analyzed data, and wrote/edited the manuscript, and C.V. designed experiments and wrote/edited the manuscript.

## Data availability

The datasets generated during and/or analyzed during the current study are available from the corresponding author on reasonable request.

**Supporting Information: Figures S1-S6**

## References

1. Scott, C. C., Vacca, F., and Gruenberg, J. (2014) Endosome maturation, transport and functions. Semin Cell Dev Biol 31, 2–10

2. Tomas, A., Futter, C. E., and Eden, E. R. (2014) EGF receptor trafficking: consequences for signaling and cancer. Trends Cell Biol 24, 26–34

3. Futter, C. E., Pearse, A., Hewlett, L. J., and Hopkins, C. R. (1996) Multivesicular endosomes containing internalized EGF-EGF receptor complexes mature and then fuse directly with lysosomes. J Cell Biol 132, 1011–1023

4. White, I. J., Bailey, L. M., Aghakhani, M. R., Moss, S. E., and Futter, C. E. (2006) EGF stimulates annexin 1-dependent inward vesiculation in a multivesicular endosome subpopulation. EMBO J 25, 1–12

5. Futter, C. E., Collinson, L. M., Backer, J. M., and Hopkins, C. R. (2001) Human VPS34 is required for internal vesicle formation within multivesicular endosomes. The Journal of cell biology 155, 1251–1264

6. Lu, Q., Hope, L. W., Brasch, M., Reinhard, C., and Cohen, S. N. (2003) TSG101 interaction with HRS mediates endosomal trafficking and receptor down-regulation. Proc Natl Acad Sci U S A 100, 7626–7631

7. Razi, M., and Futter, C. E. (2006) Distinct roles for Tsg101 and Hrs in multivesicular body formation and inward vesiculation. Mol Biol Cell 17, 3469–3483

8. Bache, K. G., Brech, A., Mehlum, A., and Stenmark, H. (2003) Hrs regulates multivesicular body formation via ESCRT recruitment to endosomes. J Cell Biol 162, 435–442

9. Henne, W. M., Buchkovich, N. J., and Emr, S. D. (2011) The ESCRT pathway. Dev Cell 21, 77–91

10. Maity, S., Caillat, C., Miguet, N., Sulbaran, G., Effantin, G., Schoehn, G., Roos, W. H., and Weissenhorn, W. (2019) VPS4 triggers constriction and cleavage of ESCRT-III helical filaments. Sci Adv 5, eaau7198

11. Adell, M. A. Y., Migliano, S. M., Upadhyayula, S., Bykov, Y. S., Sprenger, S., Pakdel, M., Vogel, G. F., Jih, G., Skillern, W., Behrouzi, R., Babst, M., Schmidt, O., Hess, M. W., Briggs, J. A., Kirchhausen, T., and Teis, D. (2017) Recruitment dynamics of ESCRT-III and Vps4 to endosomes and implications for reverse membrane budding. Elife 6

12. Adell, M. A., Vogel, G. F., Pakdel, M., Muller, M., Lindner, H., Hess, M. W., and Teis, D. (2014) Coordinated binding of Vps4 to ESCRT-III drives membrane neck constriction during MVB vesicle formation. J Cell Biol 205, 33–49

13. Davies, B. A., Azmi, I. F., Payne, J., Shestakova, A., Horazdovsky, B. F., Babst, M., and Katzmann, D. J. (2010) Coordination of substrate binding and ATP hydrolysis in Vps4-mediated ESCRT-III disassembly. Mol Biol Cell 21, 3396–3408

14. Larios, J., Mercier, V., Roux, A., and Gruenberg, J. (2020) ALIX-and ESCRT-III-dependent sorting of tetraspanins to exosomes. J Cell Biol 219

15. Babst, M., Davies, B. A., and Katzmann, D. J. (2011) Regulation of Vps4 during MVB sorting and cytokinesis. Traffic 12, 1298–1305

16. Votteler, J., and Sundquist, W. I. (2013) Virus budding and the ESCRT pathway. Cell Host Microbe 14, 232–241

17. Cho, J. H., Chang, C. J., Chen, C. Y., and Tang, T. K. (2006) Depletion of CPAP by RNAi disrupts centrosome integrity and induces multipolar spindles. Biochem Biophys Res Commun 339, 742–747

18. Kohlmaier, G., Loncarek, J., Meng, X., McEwen, B. F., Mogensen, M. M., Spektor, A., Dynlacht, B. D., Khodjakov, A., and Gonczy, P. (2009) Overly long centrioles and defective cell division upon excess of the SAS-4-related protein CPAP. Curr Biol 19, 1012–1018

19. Tang, C. J., Fu, R. H., Wu, K. S., Hsu, W. B., and Tang, T. K. (2009) CPAP is a cell-cycle regulated protein that controls centriole length. Nat Cell Biol 11, 825–831

20. Schmidt, T. I., Kleylein-Sohn, J., Westendorf, J., Le Clech, M., Lavoie, S. B., Stierhof, Y. D., and Nigg, E. A. (2009) Control of centriole length by CPAP and CP110. Curr Biol 19, 1005–1011

21. Wu, K. S., and Tang, T. K. (2012) CPAP is required for cilia formation in neuronal cells. Biol Open 1, 559–565

22. Sharma, A., Aher, A., Dynes, N. J., Frey, D., Katrukha, E. A., Jaussi, R., Grigoriev, I., Croisier, M., Kammerer, R. A., Akhmanova, A., Gonczy, P., and Steinmetz, M. O. (2016) Centriolar CPAP/SAS-4 Imparts Slow Processive Microtubule Growth. Dev Cell 37, 362–376

23. Gudi, R., Palanisamy, V., and Vasu, C. (2021) Centrosomal P4.1-associated protein (CPAP) positively regulates endocytic vesicular transport and lysosome targeting of EGFR. Sci Rep 11, 12689

24. Gudi, R., and Vasu, C. (2026) Centrosomal P4.1-associated protein is a novel regulator of ESCRT pathway function during endosome maturation. iScience 29, 114659

25. Gudi, R. R., Janakiraman, H., Howe, P. H., Palanisamy, V., and Vasu, C. (2021) Loss of CPAP causes sustained EGFR signaling and epithelial-mesenchymal transition in oral cancer. Oncotarget 12, 807–822

26. Kitagawa, D., Kohlmaier, G., Keller, D., Strnad, P., Balestra, F. R., Fluckiger, I., and Gonczy, P. (2011) Spindle positioning in human cells relies on proper centriole formation and on the microcephaly proteins CPAP and STIL. J Cell Sci 124, 3884–3893

27. Cottee, M. A., Muschalik, N., Wong, Y. L., Johnson, C. M., Johnson, S., Andreeva, A., Oegema, K., Lea, S. M., Raff, J. W., and van Breugel, M. (2013) Crystal structures of the CPAP/STIL complex reveal its role in centriole assembly and human microcephaly. Elife 2, e01071

28. Tang, C. J., Lin, S. Y., Hsu, W. B., Lin, Y. N., Wu, C. T., Lin, Y. C., Chang, C. W., Wu, K. S., and Tang, T. K. (2011) The human microcephaly protein STIL interacts with CPAP and is required for procentriole formation. EMBO J 30, 4790–4804

29. Stuchell-Brereton, M. D., Skalicky, J. J., Kieffer, C., Karren, M. A., Ghaffarian, S., and Sundquist, W. I. (2007) ESCRT-III recognition by VPS4 ATPases. Nature 449, 740–744

30. Dimaano, C., Jones, C. B., Hanono, A., Curtiss, M., and Babst, M. (2008) Ist1 regulates Vps4 localization and assembly. Mol Biol Cell 19, 465–474

31. Swift, K. A., Pustova, I., Kasberg, W., Bowman, J., Gopinath, K., Voss, E., Nelson, H., and Audhya, A. (2025) Analysis of native Ist1 dynamics reveals multiple pools of ESCRT-III on endosomes. J Cell Biol 224

32. Frankel, E. B., Shankar, R., Moresco, J. J., Yates, J. R., 3rd, Volkmann, N., and Audhya, A. (2017) Ist1 regulates ESCRT-III assembly and function during multivesicular endosome biogenesis in Caenorhabditis elegans embryos. Nat Commun 8, 1439

33. Jackson, C. E., Scruggs, B. S., Schaffer, J. E., and Hanson, P. I. (2017) Effects of Inhibiting VPS4 Support a General Role for ESCRTs in Extracellular Vesicle Biogenesis. Biophys J 113, 1342–1352

34. Cashikar, A. G., Shim, S., Roth, R., Maldazys, M. R., Heuser, J. E., and Hanson, P. I. (2014) Structure of cellular ESCRT-III spirals and their relationship to HIV budding. Elife 3

35. Harel, S., Altaras, Y., Nachmias, D., Rotem-Dai, N., Dvilansky, I., Elia, N., and Rousso, I. (2022) Analysis of individual HIV-1 budding event using fast AFM reveals a multiplexed role for VPS4. Biophys J 121, 4229–4238

36. Morita, E., Colf, L. A., Karren, M. A., Sandrin, V., Rodesch, C. K., and Sundquist, W. I. (2010) Human ESCRT-III and VPS4 proteins are required for centrosome and spindle maintenance. Proc Natl Acad Sci U S A 107, 12889–12894

37. Lin, Y. N., Wu, C. T., Lin, Y. C., Hsu, W. B., Tang, C. J., Chang, C. W., and Tang, T. K. (2013) CEP120 interacts with CPAP and positively regulates centriole elongation. J Cell Biol 202, 211–219

38. Al-Dosari, M. S., Shaheen, R., Colak, D., and Alkuraya, F. S. (2010) Novel CENPJ mutation causes Seckel syndrome. Journal of medical genetics 47, 411–414

39. Shi, L., Lin, Q., and Su, B. (2014) Human-specific hypomethylation of CENPJ, a key brain size regulator. Molecular biology and evolution 31, 594–604

40. Bond, J., Roberts, E., Springell, K., Lizarraga, S. B., Scott, S., Higgins, J., Hampshire, D. J., Morrison, E. E., Leal, G. F., Silva, E. O., Costa, S. M., Baralle, D., Raponi, M., Karbani, G., Rashid, Y., Jafri, H., Bennett, C., Corry, P., Walsh, C. A., and Woods, C. G. (2005) A centrosomal mechanism involving CDK5RAP2 and CENPJ controls brain size. Nature genetics 37, 353–355

41. McIntyre, R. E., Lakshminarasimhan Chavali, P., Ismail, O., Carragher, D. M., Sanchez-Andrade, G., Forment, J. V., Fu, B., Del Castillo Velasco-Herrera, M., Edwards, A., van der Weyden, L., Yang, F., Sanger Mouse Genetics, P., Ramirez-Solis, R., Estabel, J., Gallagher, F. A., Logan, D. W., Arends, M. J., Tsang, S. H., Mahajan, V. B., Scudamore, C. L., White, J. K., Jackson, S. P., Gergely, F., and Adams, D. J. (2012) Disruption of mouse Cenpj, a regulator of centriole biogenesis, phenocopies Seckel syndrome. PLoS Genet 8, e1003022

42. Hung, L. Y., Tang, C. J., and Tang, T. K. (2000) Protein 4.1 R-135 interacts with a novel centrosomal protein (CPAP) which is associated with the gamma-tubulin complex. Mol Cell Biol 20, 7813–7825

43. Hung, L. Y., Chen, H. L., Chang, C. W., Li, B. R., and Tang, T. K. (2004) Identification of a novel microtubule-destabilizing motif in CPAP that binds to tubulin heterodimers and inhibits microtubule assembly. Mol Biol Cell 15, 2697–2706

44. Zheng, X., Ramani, A., Soni, K., Gottardo, M., Zheng, S., Ming Gooi, L., Li, W., Feng, S., Mariappan, A., Wason, A., Widlund, P., Pozniakovsky, A., Poser, I., Deng, H., Ou, G., Riparbelli, M., Giuliano, C., Hyman, A. A., Sattler, M., Gopalakrishnan, J., and Li, H. (2016) Molecular basis for CPAP-tubulin interaction in controlling centriolar and ciliary length. Nat Commun 7, 11874

45. Breslow, D. K., and Holland, A. J. (2019) Mechanism and Regulation of Centriole and Cilium Biogenesis. Annu Rev Biochem 88, 691–724

46. Scheuring, S., Rohricht, R. A., Schoning-Burkhardt, B., Beyer, A., Muller, S., Abts, H. F., and Kohrer, K. (2001) Mammalian cells express two VPS4 proteins both of which are involved in intracellular protein trafficking. J Mol Biol 312, 469–480

47. Beyer, A., Scheuring, S., Muller, S., Mincheva, A., Lichter, P., and Kohrer, K. (2003) Comparative sequence and expression analyses of four mammalian VPS4 genes. Gene 305, 47–59

48. Dvilansky, I., Altaras, Y., Kamenetsky, N., Nachmias, D., and Elia, N. (2024) The human AAA-ATPase VPS4A isoform and its co-factor VTA1 have a unique function in regulating mammalian cytokinesis abscission. PLoS Biol 22, e3002327

49. Szymanska, E., Nowak, P., Kolmus, K., Cybulska, M., Goryca, K., Derezinska-Wolek, E., Szumera-Cieckiewicz, A., Brewinska-Olchowik, M., Grochowska, A., Piwocka, K., Prochorec-Sobieszek, M., Mikula, M., and Miaczynska, M. (2020) Synthetic lethality between VPS4A and VPS4B triggers an inflammatory response in colorectal cancer. EMBO Mol Med 12, e10812

50. Tan, J., Davies, B. A., Payne, J. A., Benson, L. M., and Katzmann, D. J. (2015) Conformational Changes in the Endosomal Sorting Complex Required for the Transport III Subunit Ist1 Lead to Distinct Modes of ATPase Vps4 Regulation. J Biol Chem 290, 30053–30065

51. Lin, H. H., Li, X., Chen, J. L., Sun, X., Cooper, F. N., Chen, Y. R., Zhang, W., Chung, Y., Li, A., Cheng, C. T., Yang, L., Deng, X., Liu, X., Yen, Y., Johnson, D. L., Shih, H. M., Yang, A., and Ann, D. K. (2012) Identification of an AAA ATPase VPS4B-dependent pathway that modulates epidermal growth factor receptor abundance and signaling during hypoxia. Mol Cell Biol 32, 1124–1138

52. Gopalakrishnan, J., Mennella, V., Blachon, S., Zhai, B., Smith, A. H., Megraw, T. L., Nicastro, D., Gygi, S. P., Agard, D. A., and Avidor-Reiss, T. (2011) Sas-4 provides a scaffold for cytoplasmic complexes and tethers them in a centrosome. Nat Commun 2, 359

53. Seluanov, A., Vaidya, A., and Gorbunova, V. (2010) Establishing primary adult fibroblast cultures from rodents. J Vis Exp

54. Kleylein-Sohn, J., Westendorf, J., Le Clech, M., Habedanck, R., Stierhof, Y. D., and Nigg, E. A. (2007) Plk4-induced centriole biogenesis in human cells. Dev Cell 13, 190–202

55. Gudi, R., Zou, C., Li, J., and Gao, Q. (2011) Centrobin-tubulin interaction is required for centriole elongation and stability. J Cell Biol 193, 711–725

56. Zou, C., Li, J., Bai, Y., Gunning, W. T., Wazer, D. E., Band, V., and Gao, Q. (2005) Centrobin: a novel daughter centriole-associated protein that is required for centriole duplication. J Cell Biol 171, 437–445

