## Supplemental Figures for "CPAP/CENPJ is essential for the stability and function of ESCRT-pathway associated AAA+ ATPase VPS4B"

Fig. S1

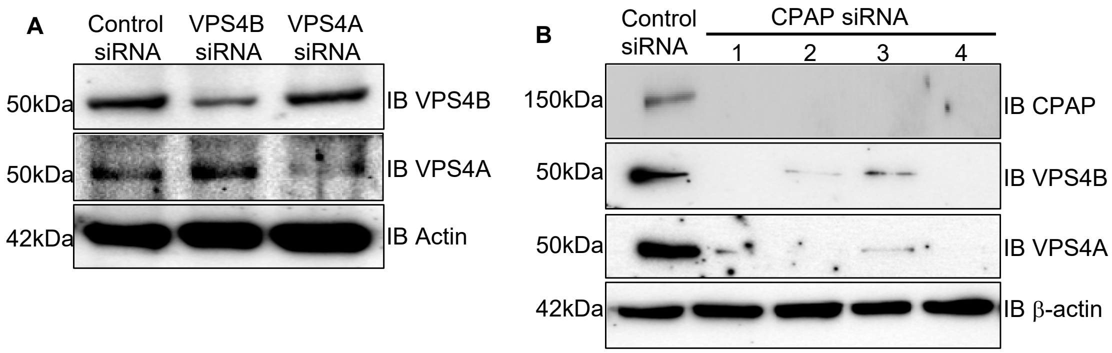


**A) Validation of VPS4 parlog-specific antibodies.** HEK293T cells were treated with control or VPS4B or VPS4A-specific siRNA for 48h and subjected to IB to detect the cellular levels of proteins using antibodies against VPS4B and VPS4B.

**B) CPAP depletion using individual CPAP-siRNAs.** HEK293T cells were treated with scrambled control or individual CPAP-specific siRNA (1: CPAP-siRNA1, 2: CPAP-siRNA2: 3: CPAP-siRNA3: and 4: CPAP-siRNA4) for 48h and subjected to IB for detection of indicated proteins.

Fig. S2

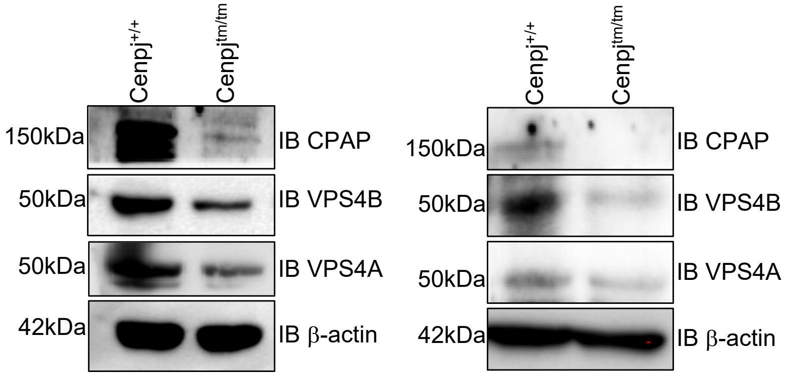


**VPS4B and VPS4A protein levels in CPAP-hypomorphic mouse cells**. Spleen cells (left panel) and primary tongue fibroblasts (right panel) prepared from 8-week-old female *Cenpj*$^{+/tm}$ (heterozygous) and *Cenpj*$^{tm/tm}$ (homozygous; hypomorphic) mice were subjected to IB for detecting indicated mouse proteins.

Fig. S3

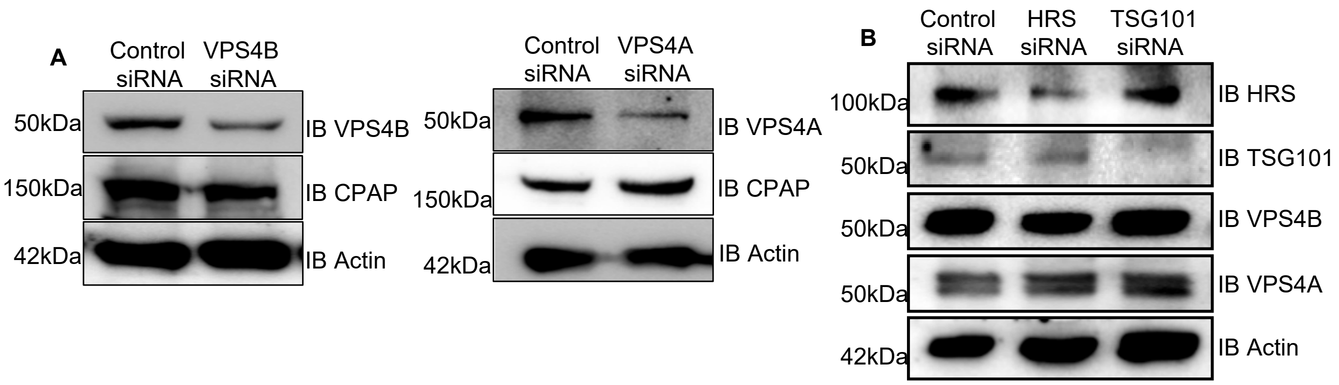


**A) Depletion of VPS4B or VPS4A proteins does not affect the cellular levels of CPAP.** HEK293T cells were treated with control or VPS4B (left panel) or VPS4A (right panel)-specific siRNA for 48h and subjected to IB to detect the indicated proteins.

**B) Depletion of early-acting protein HRS (ESCRT-0) or TSG101 (ESCRT-I) proteins does not affect the cellular levels of VPS4B and VPS4A.** HEK293T cells were treated with control or HRS or TSG101-specific siRNA for 48h and subjected to IB to detect the cellular levels of VPS4B and VPS4A proteins.

Fig. S4

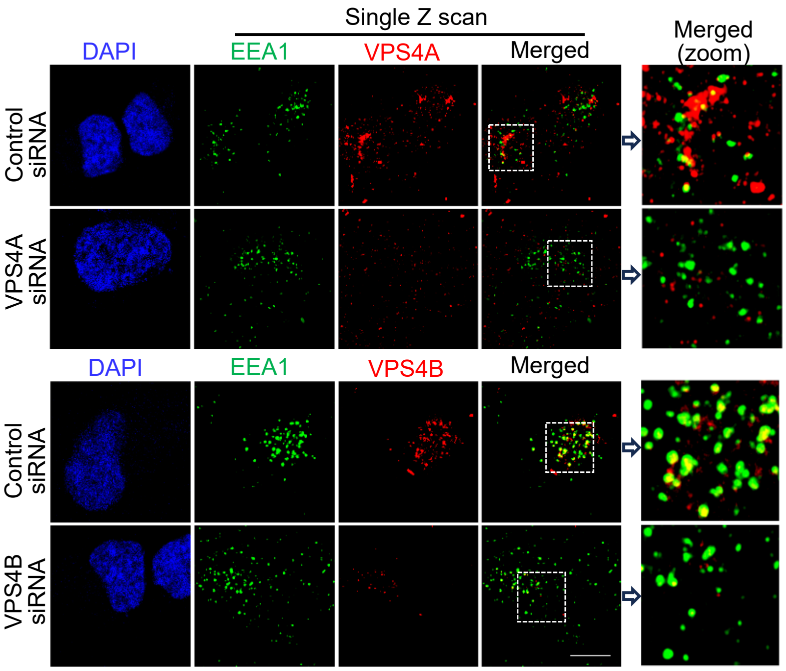


**Validating the VPS4 isoform-specific antibodies by immunofluorescence staining.** HeLa cells were treated with control or VPS4A (top panel) or VPS4B (lower panel)-specific siRNA for 48h and subjected to IB to detect their respective cellular levels using super-resolution Airyscan microscope.

Fig. S5

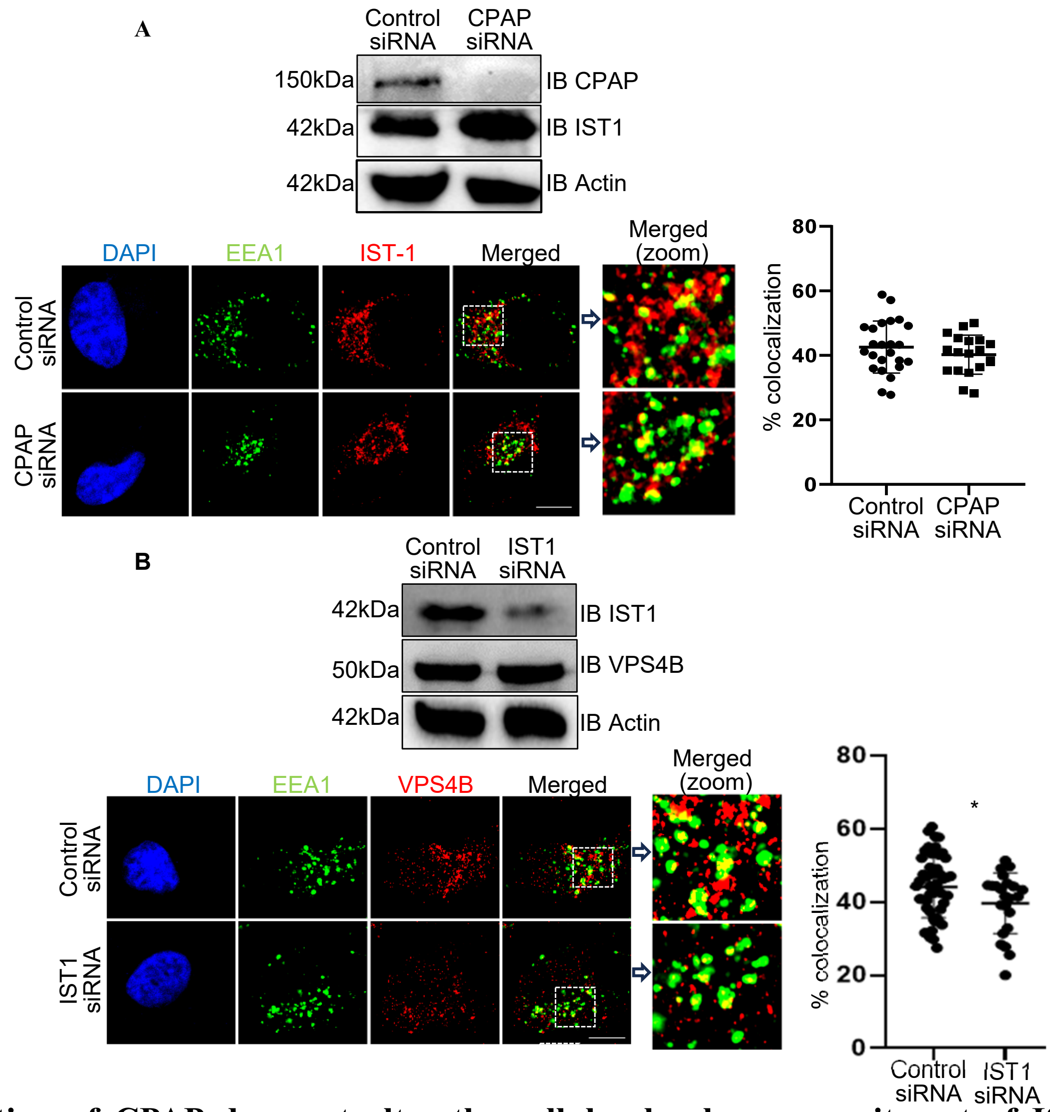


**A) Depletion of CPAP does not alter the cellular levels or recruitment of IST1 to EE.** HEK293T cells were treated with control or CPAP-siRNA for 48h and subjected to IB to detect the cellular levels of CPAP and IST1 protein (upper panel). HeLa cells expressing control or CPAP -siRNA were treated with untagged EGF for 60 min, stained for IST1 along with EEA1, and imaged by Airyscan super-resolution microscopy. Lower left panel: representative single Z-plane of images showing localization of IST1 on EEA1-positive puncta. Lower right panel: colocalization (yellow) was quantified by counting percentages of EEA1-positive (green) puncta containing IST1 protein-positive (red) puncta in representative single Z-planes of each cell and quantified from multiple cells across at least 3 experiments.

**B) Depletion of IST1 does not alter the cellular levels of VPS4B.** HEK293T cells were treated with control or IST1-siRNA for 48h and subjected to IB to detect the cellular levels of VPS4B protein. HeLa cells expressing scrambled-control or IST1 -siRNA were treated with untagged EGF for 60 min, stained for VPS4B along with EEA1 and imaged by Airyscan super-resolution microscopy. Lower left panel: representative single Z-plane of images showing localization of VPS4B on EEA1- positive puncta. Zoomed images correspond to dashed inset boxes of the indicated images. Lower right panel: colocalization (yellow) was quantified by counting percentages of EEA1-positive (green) puncta containing VPS4 protein-positive (red) puncta in representative single Z-planes of each cell and quantified from multiple cells across at least 3 experiments. Object-based colocalization macro tool of FIJI was employed. Scale bar: 10μm. p-values: *<0.05 by Mann-Whitney test.

Fig. S6

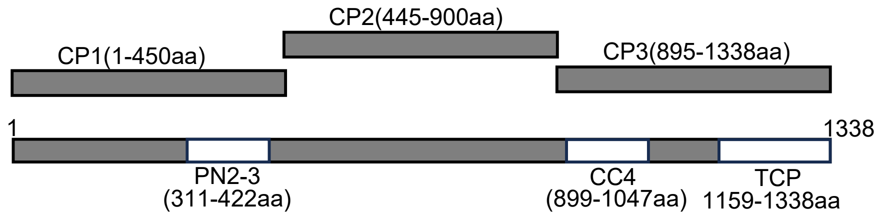


CPAP domain/fragment details, adopted from Ref # 26 (Kitagawa et al, 2011) and 37 (Lin et al 2013), are shown.
